# Processing, analysing and modelling kinetic data in the era of high-throughput single-molecule biophysics

**DOI:** 10.64898/2026.09.09.750331

**Authors:** Pim P. B. America, Misha Klein, David Dulin, Martin Depken

## Abstract

Biomolecular reactions are often composed of multiple stochastic, reversible and branched transition paths over intermediates, leading to rich dynamics. Single-molecule biophysics has revolutionized our view of biology by revealing the heterogeneity in realized paths and pointing to the importance of rare events. The recent development of high-throughput single-molecule biophysics techniques now allow to quantitatively study this heterogeneity and characterize even the rarest kinetic events. Processing, analyzing and modelling high-throughput single-molecule data has been the focus of several reports, but are often difficult to implement for non-experts. Here, we provide a guide to extract the most from transitions in single-molecule biophysics data using a first-passage time framework and maximum likelihood estimation. We specifically focused on parameter sweeps in systems with one or two characteristic timescales, and show how they can be analyzed in terms of a minimal kinetic model and its dependence on enzyme/substrate concentration, force and temperature. We introduce a general framework to perform data-driven modelling on single- and two-state models and illustrate it with concrete examples. We also provide programs with graphical user interfaces to perform such analysis on raw data, in the hope that it will empower experimental single-molecule biophysicists to extract the most out of their data.

## Introduction

Biological processes at the molecular level are stochastic (1) and are often regulated through complex, branched, kinetic pathways that can be extremely difficult to disentangle using ensemble approaches (2–5). The development of single-molecule biophysics techniques has given the tools to observe and mechanically probe such reaction, allowing us to disentangle the path-heterogeneity inherent to realizing enzymatic reactions (6–9).

Single-molecule (SM) biophysics techniques can be separated into imaging, manipulation and combinations thereof (10). The first generation of single-molecule instruments was able to monitor only a few molecules at once, limiting the depth of statistical analysis that could be performed. In recent years, several technological breakthroughs have significantly improved size and quality of data sets produced. The introduction of cameras with larger field of views and higher frame rates, such as complementary metal-oxide-semiconductor (CMOS), dramatically increased the throughput of camera-based assays. Furthermore, the parallelization capabilities of graphics processing units (GPUs) have sped up position tracking algorithms to monitor hundreds to thousands of molecules in parallel and in real-time (11–13). Altogether, these developments have lead to the era of high-throughput single-molecule biophysics, both for single-molecule fluorescence (14, 15) and force spectroscopy assays (16–19). Throughput is now also sufficiently large for single-molecule experiments to be combined with next-generation sequencing techniques to explore sequence specific effects (11, 13).

The increasing size and complexity of single-molecule datasets have driven the need for more advanced methods for data processing, analysis, and kinetic modelling. Improved statistical resolution reveals increasingly complex reaction dynamics, uncovering kinetic states and pathways that were previously inaccessible (20). Although substantial progress has been made in modelling the macromolecular dynamics underlying biological processes, many earlier studies were restricted by the amount of data obtained (21–27). Single-molecule assays provide direct access to molecular states and the transition times between them, enabling the reconstruction of detailed kinetic schemes connecting these states (9, 28–32). However, not all molecular states that influence the observed kinetics are experimentally resolvable. Limited spatiotemporal resolution can obscure transient or short-lived intermediates, resulting in hidden molecular states and transition pathways that remain inaccessible to direct observation (33). Nevertheless, these hidden states and their associated transition rates might still be inferred by combining dwell-time analysis across a range of experimental conditions with kinetic modelling (18, 34–38).

Here, we provide a step-by-step methodology describing how to process and analyze single-molecule data to explain it in the context of a minimal model. We first review several common single-molecule approaches and the two classes of data they typically yield. We cover step-like activity traces with clearly defined plateaus, such as obtained from various single-molecule experiments (32, 34, 36, 39–41). Next, we cover continuous activity traces with processive directionality, such as obtained from translocating motors using magnetic tweezers, optical tweezers or flow-stretch experiments (17, 20, 42). We describe how the activity traces collected under varying experimental conditions can be processed into dwell-time distributions, and how modelling and maximum likelihood estimation (MLE) can be used to determine the underlying reaction scheme. Moreover, we provide the methods to perform data-driven model selection based on single-molecule data with high statistics. To exemplify these methods, we cover systems with up to two timescales and show how to constrain the minimal model that explains the data. We illustrate the power of this approach through practical examples to help the interested reader with extracting the most information from their own data. Furthermore, we provide GUIs to process the single-molecule traces into a dwell-time distribution and perform MLE fitting for complex distributions.

### Extracting empirical dwell-time distributions from single-molecule traces

Various assays can be used to investigate interactions at the single-molecule level, such as single-molecule force spectroscopy and fluorescence (7). Here, we focus on techniques that monitor enzymatic activity over long periods of time, such as a nucleic acid or protein dynamically changing conformational states (**Figure 1AB**), and molecular motors processing nucleic acids **(Figure 1CD**). These experiments typically monitor the position of a trapped bead or the fluorescence intensity as a function of time, resulting in the two archetypical time-traces we present in **Figure 1EF**.

**Figure 1:**
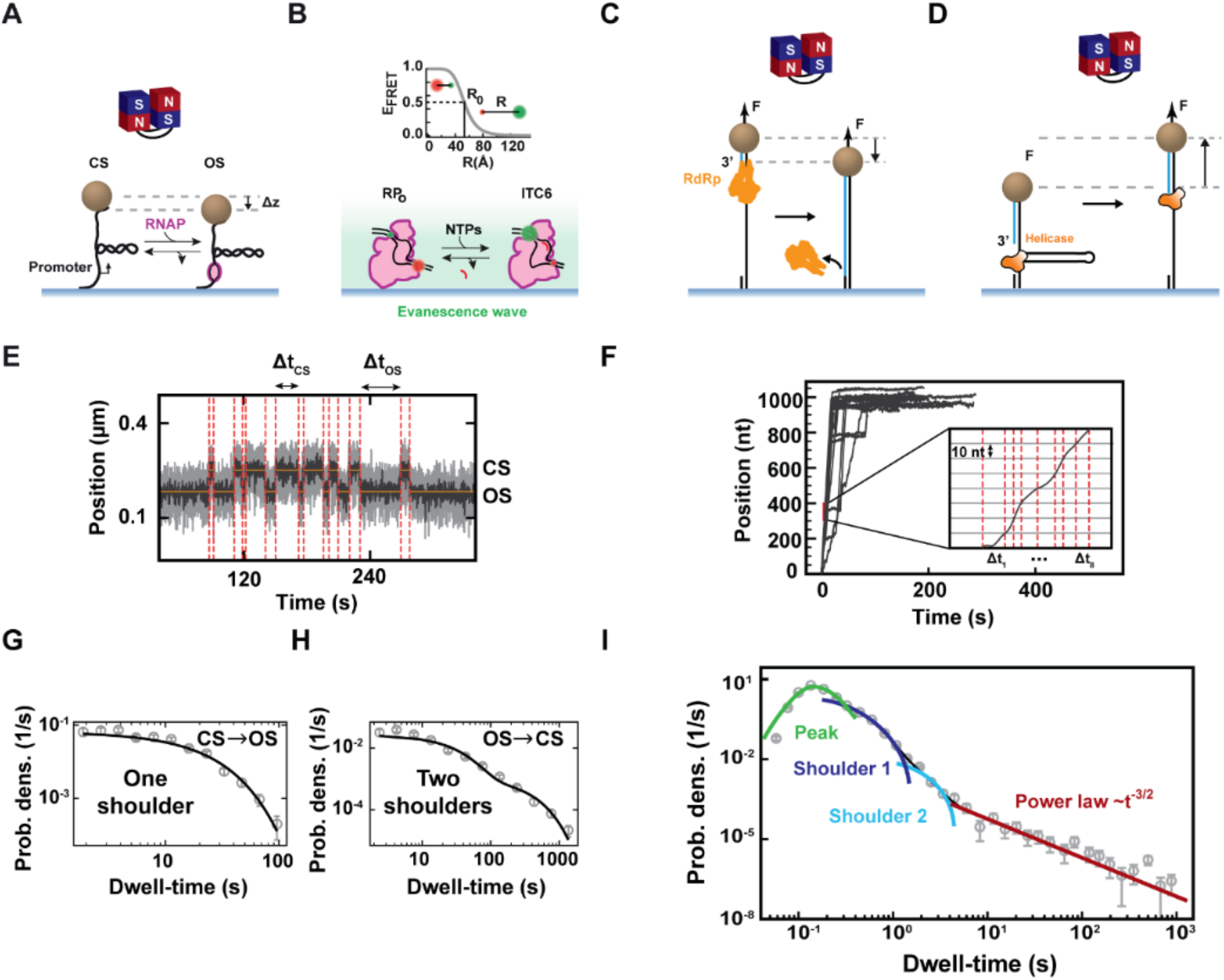
Examples of single-molecule biophysics experiments with the typical time-traces and the applied dwell-time analysis. **(A)** Schematic of a magnetic tweezers assay to study open-complex formation by bacterial RNA polymerase (RNAP) on its DNA promoter. The transition from the closed state (CS) to the open state (OS) was measured from the increase in supercoils in the DNA tether, decreasing the bead height by Δ*z*. **(B)** Schematic of a FRET assay to monitor the dynamics of bacterial transcription initiation. The donor dye is upstream on the non-template DNA, while the acceptor dye is located downstream on the template DNA. In the presence of nucleotides, the RNAP scrunches the DNA, changing the donor-acceptor distance and therefore the FRET efficiency. **(C, D)** Schematics of magnetic tweezers assays to investigate either the elongation dynamics of the RNA dependent RNA polymerase (RdRp) on a single-stranded RNA template (C) or double-stranded nucleic acid unwinding dynamics by a helicase (D). **(E)** Example time-trace obtained in a magnetic tweezers experiment described in (A) showing transitions between CS and OS with timescales Δ*t*_CS_ and Δ*t*_OS_. **(F)** Example SARS-CoV-2 RTC elongation traces for the experiment described in (C). Inset: zoom-in of the trace with red dashed lines indicating when the 10 nt dwell-time window is crossed. **(G, H)** Dwell-time distributions obtained for the CS-to-OS transitions of the RNAP-promoter complex (G) and for the OS-to-CS transitions (H). **(I)** The dwell-time distribution for the traces represented in (F) using a 10 nt dwell-time window (grey circles) and the corresponding fit-function consisting of a peak (gamma distribution), two shoulders (exponential distributions) and a power law decay (∼*t*^−3/2^). The error bars on the bins are one standard deviation from 1000 bootstraps. (A, E, G, H) are adapted from (39). (B) is adapted from (40). (C, F, I) are adapted from (29).

First, we have single-molecule measurements on a nucleic acid or protein dynamically changing conformational states. As an example, we consider magnetic tweezers and single-molecule FRET experiments monitoring the bacterial RNA polymerase (RNAP)-promoter open complex (RP_O_) dynamics (43, 44) (**Figure 1AB**). In the magnetic tweezers assay, a negative or positive supercoil is induced in a DNA molecule by rotating the magnets located above the flow chamber and thus the magnetic bead, resulting in plectoneme formation (45). For the positively supercoiled case, the opening of the promoter by the RNAP induces the addition of approximately one loop to the plectoneme, decreasing the measured extension of the DNA tether by Δ*z* in **Figure 1A** (39). The opening and closing of the RNAP-DNA complex appear as sudden increases and decreases in the magnetic bead’s height. Detection of the transitions between open and closed RNAP-DNA complexes was performed using a change point analysis to detect these clearly defined steps (**Figure 1E**) (39, 46). The collected ‘dwell-times’ for either the closed (CS) or open state (OS), Δ*t*_CS_ and Δ*t*_OS_ respectively, where assembled into distributions (39) (**Figure 1GH**).

Using single-molecule FRET, the RP_O_ dynamics and the dynamics of initial transcription can be monitored as changes in the FRET efficiency (40, 43, 47), providing that the dyes have been positioned appropriately to maximize the FRET efficiency difference (**Figure 1B**) (14). Hidden Markov modelling is often used to capture the probabilities for the transitions between states with a distinct FRET efficiency representing conformational states of the biomolecules (48). However, the hidden states in Hidden Markov models are typically not interpretable as physical states (49, 50). To identify physical states that are indistinguishable in terms of their FRET efficiencies, it might be possible to apply the data-driven model selection procedure described in this manuscript.

For both the magnetic tweezers (**Figure 1E**) and FRET (40) assay, clear discrete transitions between molecular states were observed, which can be detected using different step finder algorithms (46, 50–52). The choice of detection algorithm depends on the spatiotemporal resolution of the measurement, the number of molecular states of interest, transition times on widely different timescales (48), and the signal fluctuations (53). The life-times in each measured molecular state can be directly obtained (**Figure 1E**) and assembled into empirical dwell-time distributions (**Figure 1GH**) (54). Characteristic timescales and probabilities can be fitted out from the distributions and compatible models and their parameters can be inferred.

The second major class of single-molecule measurements relates to processive molecular motors converting nucleic acids, DNA or RNA, from double-stranded to single-stranded or the reverse. For example, the traces from a magnetic tweezers assay monitoring the nucleotide addition cycle of a viral RNA polymerase on ssRNA (**Figure 1C**) or a viral helicase opening an RNA hairpin (**Figure 1D**). The traces of this motor activity display continuous transitions through molecular positions (**Figure 1F**). Molecular motors are typically nanometer sized and perform steps of dimensions defined by their substrate: RNA polymerases move by single DNA base pairs (∼0.34 nm) every 20-50 ms while a ribosome moves by codon of three nucleotides (∼1.2 nm) every 100-200 ms (8). Therefore monitoring single translocations of these motors requires a low experimental noise to see each step within the time window (55). When the spatiotemporal resolution is sufficient to observe discrete steps by the motor, the traces classify as the first type of single-molecule traces and dwell-time analysis can be performed as described above (30, 31, 33, 56). When the typical size and duration of the steps fall below the spatiotemporal resolution of the instrument, a continuous time-trace is observed, as for our magnetic tweezers assay on the nucleotide addition cycle of viral polymerases (**Figure 1F**) (18, 29). To extract more information from these traces, a coarse-grained dwell-time analysis can be performed by dividing the traces into non-overlapping windows larger than the spatiotemporal resolution. When a large number of full-length continuous time-traces is recorded, the large statistics enables dwell-time analysis (**Figure 1I**) and the underlying kinetic schemes of molecular states can be unravelled (18, 29, 57).

### Interpreting dwell-time distributions

In the context of SM experiments, dwell-times represent the time it takes to exit a state, also referred to as the first-passage time (FPT) from entry into to exit out of that state (54). Dwell-times originating from a single state are exponentially distributed (**Methods**) and as a result dwell-times originating from exits through one or multiple internal states typically give rise to multi-exponential distributions. For simplicity and clarity, we will only discuss distributions that have at most two timescales and the minimal models describing them. The methodology can be generalized to more complex reaction schemes, as will be touched upon in later chapters.

First-passage time distributions with one and two timescales can be captured by a single- and double-exponential probability density function (pdf) respectively

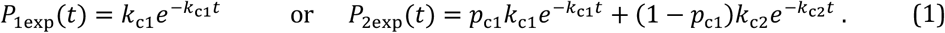

Here, *k*_c*i*_ with *i* = 1, 2 are the characteristic rates of the first and second exponential and *p*_c*i*_ are the probabilistic weights that must add up to one to ensure normalization. When a single timescale dominates the reaction, the resulting dwell-time distribution follows a single exponential decay, which appears as a single “shoulder” on a log–log scale (**Figure 2A**). In reactions involving two characteristic timescales, two distinct types of distributions can result depending on the underlying process. In the first type, both weights are positive, resulting in a distribution with two distinct “shoulders” on a log-log scale when the timescales are well-separated. In the second type, one weight is negative, producing a peaked distribution characterized by a linear rise followed by an exponential decay (**Figure 2A**).

**Figure 2:**
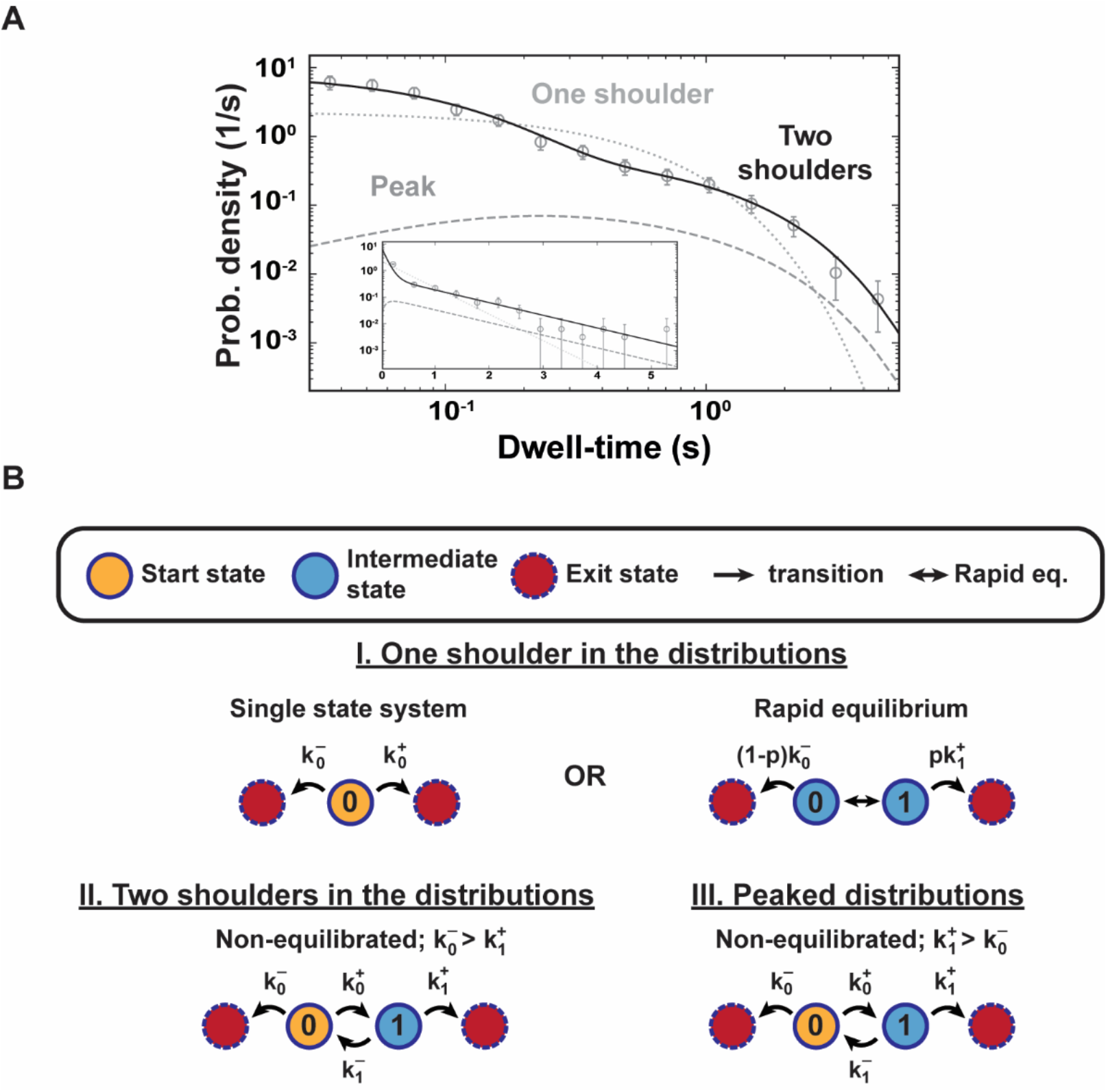
Types of dwell-time distributions for different two-state systems. **(A)** A single and a double exponential dwell-time distribution on log-log representation, appear as either one (dotted curve) or two shoulders (full curve) respectively in a log-log representation. A (pseudo-) gamma dwell-time distribution (an exponential rise followed by an exponential decay, dashed curve) is peaked. (inset) The single exponential, double exponential and pseudo-gamma distributions on log-linear representation. The grey circles with error bars represent the mean bin values and one standard error of the mean (SEM) drawn for a generated double exponential dwell-time distribution of 1000 datapoints. **(B)** Two-state models grouped per dwell-time distribution type (I-III). The start state is indicated as a yellow disc, the intermediate state in blue and the exit states in red. Kinetic steps are indicated as arrows between states. A rapid equilibrium is indicated as a double arrow.

Dwell-time distributions are also commonly represented on a log-linear scale. In this representation, each exponential decay appears as a linear region. However, as illustrated by the example containing two exponential decays in **Figure 2A**, one of the linear regions may become extremely short when the timescales are well-separated and may therefore be represented by only a single bin. For this reason, we recommend the use of a log–log representation, since features associated with processes occurring on different timescales are more clearly distinguished.

Importantly, the type of dwell-time distribution observed teaches us about the minimal underlying system. In **Figure 2B**, the single- and two-state models are grouped according to their corresponding dwell-time distribution types, that is single- or double exponential or peaked (**Figure 2A**). The figure presents all possible reaction schemes containing up to two states, represented by one or two characteristic timescales, and identifies the minimal model capable of describing the dwell-time distributions shown in **Figure 2A**. In these schemes, states are represented as discs connected by transition rates, indicated by arrows. The dwell-time is defined as the time required to transition from the initial state (state 0, yellow disc) to an exit state (red discs), either directly or after multiple cycles involving transitions to and from an intermediate state (state 1, blue disc) (**Figure 2B**). Now we will describe what we learn from the distribution types about the underlying systems.

The observation of a single shoulder in the dwell-time distribution indicates that the underlying reaction scheme can be summarized by a single-state system with two exit rates or an equilibrated two-state system, which is also an effective single-state model (**Figure 2B**) as long as the equilibration time is below the dynamic range of the experiment. In either case the characteristic rate is expressed as

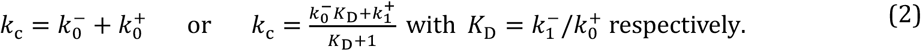

The shoulder is then fitted with a single exponential distribution with characteristic rate *k*_c_ (**Equation 2**).

If the dwell-times exhibit two shoulders or a peak, the minimal model that could capture the data is an out-of-equilibrium two-state system. When 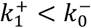 such a system produces a double-shoulder distribution, while for 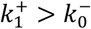 it produces a peaked distribution (**Figure 2B**), as derived in the **Methods**. In both cases, the characteristic rates *k*_c1_, *k*_c2_ and probability *p*_c1_ can be expressed in terms of the microscopic rates as

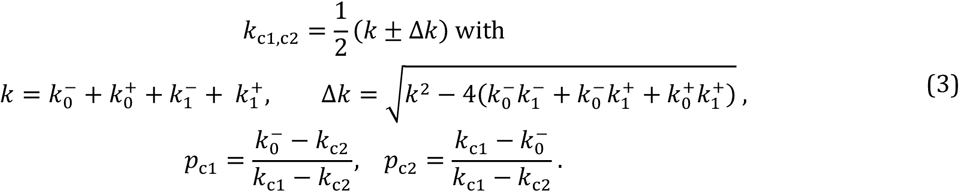

However, since a double exponential distribution has only three independent parameters *k*_c1_, *k*_c2_ and *p*_c1_, the four microscopic rates from the two-state system cannot be directly determined from fitting a single distribution. To distinguish between the various models that give the same distribution type, we need to access the change in the dwell-time distributions when modulating at least one microscopic rate in the system.

### Sweeping rates by sweeping experimental conditions

Reaction rates can depend on various parameters, such as substrate or enzyme concentration, a mechanical force applied to the system or the temperature.

#### Concentration

For example, the binding rate of enzyme *E* at concentration *C* to a single substrate in solution, such as RNA polymerase binding from solution to a single DNA promoter (**Figure 1A**), follows first-order kinetics and therefore depends linearly on the enzyme concentration

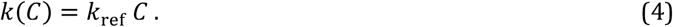

When the enzyme concentration *C* = [*E*] is expressed in nanomolar (nM), which is typical for single-molecule experiments (1), the parameter *k*_ref_ corresponds to the binding rate at an enzyme concentration of 1 nM. In the situation, where substrate molecules from solution bind to a single enzyme, the enzyme concentration is replaced by the substrate concentration *C* = [*S*].

#### Force

Mechanical forces applied to the reactants can also modulate reaction rates by altering the free-energy of a reactions transition state. This is particularly relevant when the reaction requires a conformational change in the enzyme or substrate working with/against an external tension *F* that is applied, e.g. to the DNA tether in **Figure 1A**. During such conformational transitions, the extra mechanical work *F* ⋅ *dx* needs to be performed to reach the transition state situated distance *dx* away along the direction of the applied force. This distance is typically measured in nanometers, and the resulting force dependence of the reaction rate is commonly described by an Arrhenius-type relation (21)

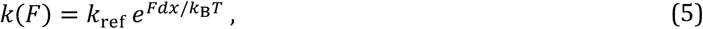

in which *k*_ref_ represents the rate at zero force. In single-molecule experiments, the applied force *F* is generally measured in piconewtons (pN). In practice, however, identifying which transition rates in a kinetic scheme exhibits a force dependence is often nontrivial.

#### Temperature

Every reaction rate has a temperature dependence that follows the Arrhenius law, characterized by an activation energy *E*_act_ (1)

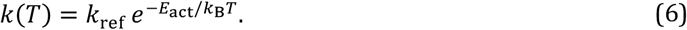

In this description, *k*_ref_ corresponds to the rate in the limit of very high temperature (*k*_*B*_*T* ≫ *E*_act_). Because this dependence is exponential, even small fluctuations in temperature can produce substantial variations in the measured reaction rates when *E*_act_ ≫ *k*_*B*_*T*. This highlights the importance of precise temperature control during single-molecule experiments (58).

As discussed above, the dwell-time distribution of a single- or two-state system measured under a single condition does not contain sufficient information to uniquely determine all microscopic rates from a fit to a single or two exponential pdf, respectively (**Figure 2, Equations 2 and 3**). In principle, one could identify the correct model by globally fitting all possible single- and two-state systems, each containing a single concentration- or force-dependent rate, to a series of dwell-time distributions measured across a parameter sweep.

In practice, however, this approach presents an important ambiguity: a poor fit may arise either because the proposed model is incorrect or because the fitting algorithm failed to converge to the global optimum. To address this ambiguity, we instead propose a systematic procedure to identify the minimal model capable of describing the data based on the characteristic changes in the dwell-time distributions as a function of enzyme or substrate concentration, externally applied force acting on a single rate or temperature.

### Guidelines for model selection based on dwell-time distributions

In this section, we describe our recommended procedure for systematically selecting a minimal model that describes single-molecule dwell-time data. We distinguish between two approaches: data-driven model selection and data-oriented model selection. In the data-driven approach, the minimal model is inferred directly from the empirical dwell-time distributions without requiring prior knowledge of the underlying reaction mechanism. Prior knowledge is used only afterwards to interpret and validate the resulting model. In contrast, data-oriented model selection incorporates existing knowledge of the reaction mechanism to construct a single minimal model consistent with both the data and the known biochemical or biophysical constraints.

(Step 1) Determine the minimal number of effective states and microscopic rates in the model.

a. Determine the maximal number of timescales in the dwell-time distributions in sweeps of enzyme/substrate concentration, force or temperature, which determine the minimal number of effective states (Data-driven model selection).
b. Find the rate dependencies in the system and whether microscopic rates can be removed based on various limits of the sweeps (Data-driven model selection).
c. When multiple models with the same number of rates are consistent with the data, turn to knowledge from the literature to determine a single model that describes the data (Data-oriented model selection).

(Step 2) Build the model and perform the model fit to the data to estimate the microscopic rates.

(Step 3) Evaluate the fitted model parameters, make a biological/chemical interpretation of the model based on the literature and predict unused data/literature with the fitted model for validation.

In the following, we discuss the considerations relevant to each step of the procedure in general and in the context of single- and two-state systems with a single concentration-dependent rate. We then illustrate the data-driven and data-oriented model selection procedures using example datasets, demonstrating how the combined analysis of concentration dependencies and temperature can reveal a hidden state, and how the minimal kinetic model can be identified when no rate dependency is observed.

#### Step 1a

To determine the number of characteristic timescales in the dwell-time distributions for the various conditions measured, inspect the distributions in a log-log representation and identify clearly distinguishable features such as shoulders or peaks (**Figure 2A**). The confidence intervals of the histogram counts on log-log scale, represented by the error bars on the bins, can be estimated by repeatedly resampling the dwell-times, for example using bootstrap resampling with 100-1000 iterations (59). The observed features in the distributions should subsequently be quantified by fitting the dwell-times. The number of distinct timescales observed in the distributions determines the minimal number of effective states required in the kinetic model.

As discussed above, for all limiting cases of a single and two-state system, the dwell-time distributions present as a single shoulder, two shoulders or a peak on log-log scale (**Figure 2**). When the distributions for the concentration sweep are all a single shoulder and well-fitted with a single exponential distribution, the system is defined by a single characteristic timescale or rate and can be represented by either a single-state model or an equilibrated two-state model. The peaked distribution is determined by two timescales, one for the rise and the other for the decay. Two shoulders in the distribution are also defined by two inherent timescales. For both distribution types, the minimal model is a non-equilibrated two-state system (**Figure 2**).

#### Step 1b

To determine the rate dependencies in the system and whether the model can be further simplified, we investigate the dwell-time distributions over the full range of the dependency: from the limit in which the modulated rate is very small compared to the other rates to the limit of a very large rate compared to the other rates. The procedure to determine the minimal model for a two-state system with a single rate dependency on concentration *C* is illustrated in **Figure 3**.

**Figure 3:**
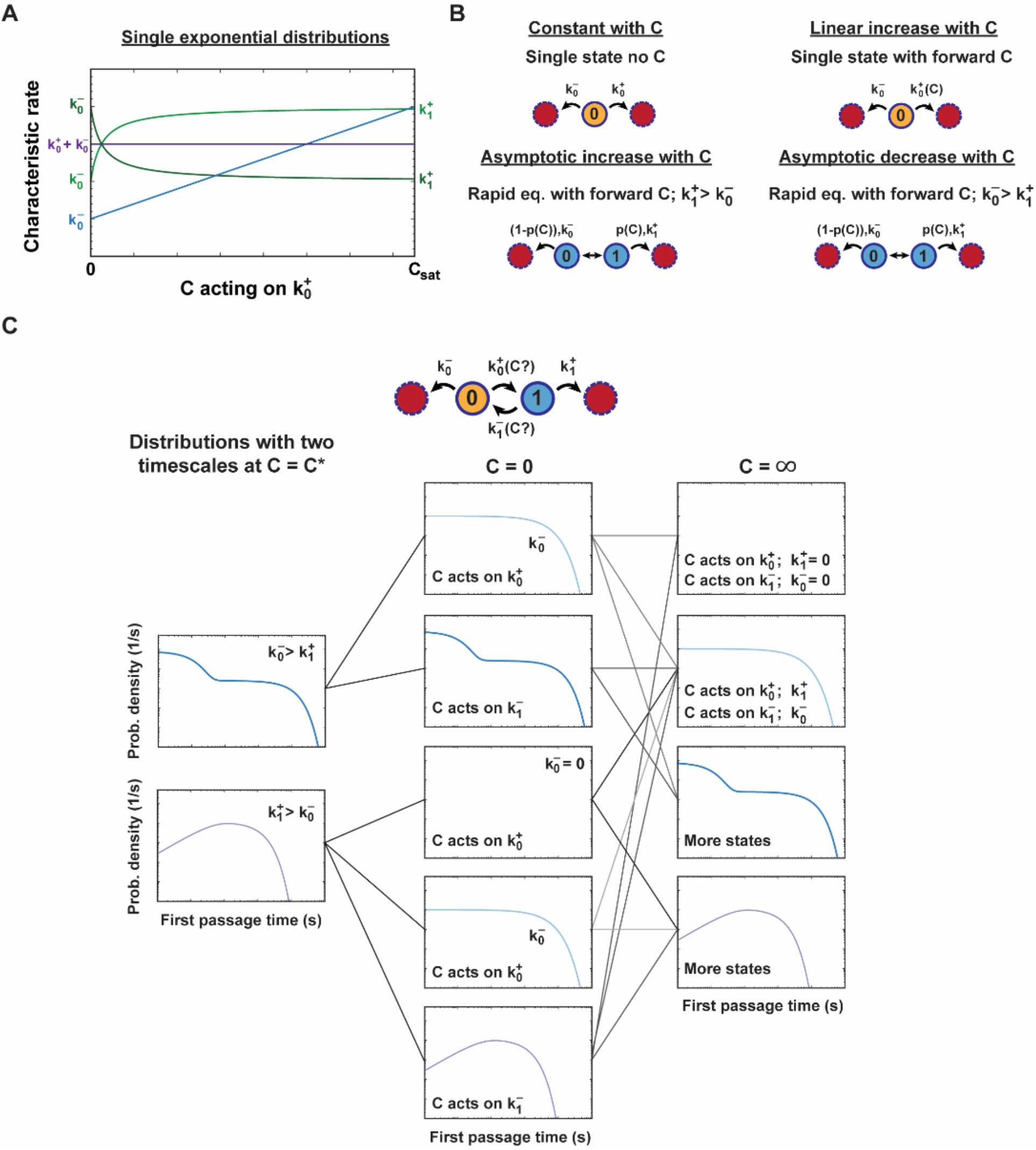
Method to parameterize single- and two-state systems based on the dwell-time distributions with a single rate dependency on concentration *C*. **(A)** The typical trends for single- or two-state models in the characteristic rate with a *C* dependency on the internal forward rate 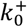. The characteristic rate is either constant (purple line), increases linearly with *C* (blue line) or increases or decreases asymptotically with *C* (light or dark green line). **(B)** The minimal models in case of single shoulder dwell-time distributions per trend in the characteristic rate with *C*. **(C)** Flow diagram to parameterize the non-equilibrated two-state models with a *C* dependency on either the internal forward rate 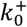 or the backward rate 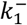 (top) based on the dwell-time distribution types at three conditions. i.e. at some intermediate *C*^∗^, *C* = 0 and very high *C*(= ∞). The model selection procedure is explained in section “**Guidelines for model selection based on dwell-time distributions**”.

When a single shoulder describes the dwell-time distributions for all measured values of *C* and are well-fitted with a single exponential distribution (**Equation 1**), the system is described either by a single-state model or by an equilibrated two-state model, as discussed above. These cases can be distinguished by analysing the dependence of the fitted characteristic rate on *C* (**Figure 3AB**). If the characteristic rate is independent of *C*, the system corresponds to a single-state process without concentration dependence. A linear increase in the characteristic rate with *C* indicates a concentration-dependent transition rate, while a nonzero characteristic rate at *C* = 0 implies the presence of an additional concentration-independent exit pathway entered with probability 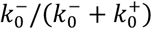 and transition rate 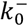 (**Figure 3B**). In contrast, an asymptotic increase or decrease in the characteristic rate with *C* results from a two-state system, for which the internal transition rates have a concentration dependence and can be assumed in rapid equilibrium (**Methods**).

In case the dwell-time distributions in the intermediate regime of *C*(= *C*^∗^) exclusively present two shoulders or a peak, fitted with either the sum of or the difference between two exponential distributions respectively (**Equation 1**), the system must instead have two non-equilibrated states and the *C* dependence is on the internal forward or backward rate 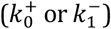 (**Figure 3C**). The type of distribution directly reVlects the relative magnitudes of the exit rates: a double exponential distribution indicates 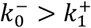, whereas a peaked distribution indicates 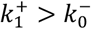 (**Figure 2B, Figure 3C**). When the distribution type is unchanged at *C* = 0, *C* is acting on 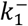. Alternatively, when the distribution type is a single shoulder or it is empty at *C* = 0, *C* is acting on 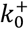 and the characteristic rate sets 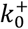. For an empty distribution in this limit, we have 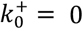. To check whether exit rates can be set to zero in all other cases and if the system can be fully described by a two-state system, the dwell-time distribution should be obtained at very high *C*(→ ∞). When the distribution becomes a single shoulder in this limit, the asymptotic rate represents the exit rate in the direction of the *C* dependency (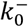 or 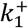). In case the events take much longer than biologically relevant timescales at very high *C*, the dwell-time distribution is considered empty and the exit rate in the direction of the *C* dependency can be set to zero. When two timescales are distinguished from the distribution in the limit of very high *C* (i.e. two shoulders or a peak), more complex kinetics applies than a two-state system (**Figure 3C**).

To complete the classification of non-equilibrated two-state systems with a single rate dependence on *C*, consider that *C* could also modulate one of the exit rates (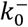 or 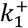) (**Figure S1**). In these systems, the dwell-time distributions continuously transform as *C* changes, unlike the models with an internal microscopic rate dependence, where double exponential and peaked distributions are mutually exclusive. For example, when 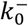 depends on *C*, the distributions transform from peaked to single exponential, then to double exponential and eventually back to single exponential (**Figure S1B**). When 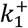 depends on *C*, the distributions transform from peaked to single exponential, to double exponential and back to single exponential (**Figure S1C**). Together, these criteria allow all two-state systems to be distinguished.

#### Step 1c

If multiple models remain consistent with the data, additional prior knowledge about the reaction mechanism is required to identify a unique minimal model. Relevant questions include: Which intermediates have been experimentally observed? Which transition rates are known to depend on enzyme or substrate concentration? Are certain rates significantly faster or slower than the other relevant transitions? To address these questions, we recommend integrating kinetic information obtained from bulk and single-molecule experiments.

#### Step 2

To fit the model to the dwell-times for each condition measured, we recommend using a maximum likelihood estimation (MLE) approach, which fits the individual dwell-times directly and therefore avoids binning-related artifacts (60).

#### Step 3

When evaluating the fitted model parameters and predicting data that was not used for model selection, it is important to recognize that the experimentally accessible range of dwell-times is limited by factors such as the acquisition frequency, spatiotemporal resolution, total acquisition time, limited statistics, and the lifetime of fluorescent probes. Consequently, the inferred model is only validated within the experimentally observable time range.

To make a biological/biochemical interpretation of the model, we encourage to combine kinetic information from bulk and single-molecule experiments with other biological assays.

### Modulating force and temperature dependencies

Similar to the example procedure with a single rate dependency on concentration (Step 1a and 1b), the limiting cases of two-state systems could be determined from a force dependency on one of the microscopic rates or from the temperature dependency.

When there is a force dependence on a single rate (**Equation 5**), the limiting case of the two-state system can be determined by modulating the force across a range from strongly opposing conditions, where the microscopic rate approaches zero (analogous to *C* = 0, **Figure 3**), to strongly assisting conditions, where the microscopic rate becomes very large (analogous to *C* → ∞, **Figure 3**). The trends in the characteristic rate with force will, however, be different since reaction rates have an exponential dependence on the modulated force (**Figure S2, Equation 5**), while a dependency on concentration is linear (**Equation 4**).

In practice, however, some experimental setups cannot access the full force range from opposing to assisting conditions. As a result, certain regions of parameter space may remain inaccessible within a single assay, preventing complete characterization of the system. In such cases, complementary assays probing different regions of parameter space can be combined (29, 61). A force could also act on both the forward and backward transition rate, making it more complicated to disentangle the rate dependencies.

The temperature acts on every microscopic rate (**Equation 6**), but in specific cases a characteristic trend of the underlying states can be found in the shifts in the dwell-time distribution, e.g. a bimodality in the characteristic rate with an equilibrated second state (**Figure S3**).

### Example study: RNAP-promoter open-complex formation

To illustrate how the data-driven model selection works, we use magnetic tweezers data from (39) describing the dynamics of the bacterial RNAP open complex formation on the DNA promoter. We show how the model for the open-complex formation of bacterial RNA polymerase (RNAP) on a promoter could be obtained from an RNAP concentration and temperature sweep (39), when the information on the reaction from literature is only used to interpret and validate the model. Then we exemplify the data-oriented model selection by elaborating on how the model for the closing of the complex was determined without rate dependency (39).

#### CS-to-OS transitions: Kinetic model for the RNAP-promoter open-complex formation

Here, we recapitulate how the underlying kinetic model for the transition from the closed complex (CS) to the open complex (OS) could be inferred from the dwell-time distributions and their dependence on RNAP concentration and temperature, following the data-driven model selection procedure (no Step 1c).

##### Step 1a

For the transitions from CS to OS, the dwell-time distributions exhibited a single shoulder at all measured RNAP concentrations and temperatures, as in **Figure 1G** (39). These distributions were well-described by a single exponential pdf defined by a single characteristic timescale, indicating that the transition can be represented either by a single-state system or by a two-state system in rapid equilibrium (**Figure 2**).

##### Step 1b

To determine the minimum number of effective states and microscopic rates in the system and the rate dependencies, we analysed an RNAP concentration (i) and temperature (ii) sweep of the effective CS-to-OS transition rate *k*_open_ (empirical characteristic rate).

i. RNAP concentration sweep: in the effective CS-to-OS transition rate *k*_open_ with RNAP concentration a close to linear increase is observed, starting from zero (**Figure 4A**), indicating that there is a linear RNAP concentration dependency on one microscopic forward rate, such as **Equation 4**. Since there is no open-complex formation at zero concentration (**Figure 4A**), we conclude that there is no exit from the starting state 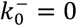. The measured concentration range was, however, insufficient to determine whether the system is a single-state or an equilibrated two-state system.
ii. Temperature sweep: the effective CS-to-OS transition rate *k*_open_ displayed a bimodal behaviour with temperature, which acts on all microscopic rates (**Equation 6**). The rate increased linearly on a logarithmic scale between low and intermediate temperatures (25–37 °C) and decreased again at higher temperatures (37–45 °C) (**Figure 4B**). This non-monotonic temperature dependence demonstrates that the CS-to-OS transition cannot be described by a single-state process and instead requires at least an effective two-state system (**Figure 4B, Figure S3**).

**Figure 4:**
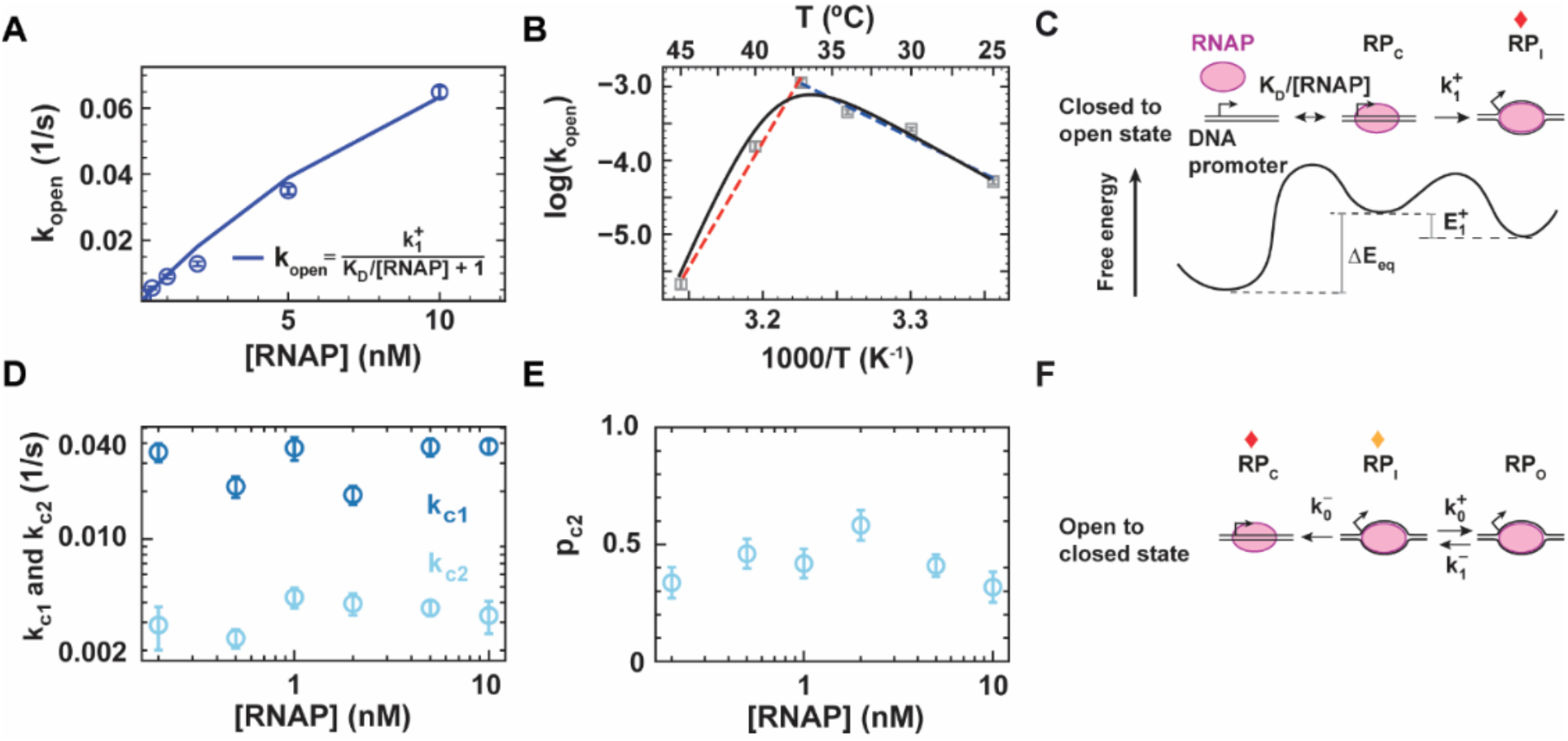
Kinetic modelling of RNAP-DNA promoter open-complex dynamics observed in single-molecule experiments. **(A)** The rates of RNAP-DNA promoter open-complex (OS) formation *k*_open_ versus RNAP concentration (nM) obtained from the single exponential fits to the dwell-time distributions for open-complex formation (CS → OS) are shown as circles with error bars, representing the mean and one standard error of the mean (SEM) from 1000 bootstraps. The fit of the model for RNAP-promoter open-complex formation is shown as a solid line. **(B)** Temperature dependence of the RNAP-DNA promoter open-complex formation rate *k*_open_. **(C)** Kinetic model describing the opening of the RNAP-DNA promoter complex and its free energy landscape. The red diamond indicates the end of the FPT. **(D)** Fast and slow exponential rates (*k*_c1_ and *k*_c2_) and **(E)** probability of the slow exponential *p*_c2_ obtained from the double exponential fits to the dwell-time distributions of the open to close complex transition (OS → CS) (Fig. 1E) as a function of RNAP concentration (nM). The circles with error bars represent the mean rate values and one SEM from 1000 bootstraps. **(F)** Kinetic model of the closing of the RNAP-DNA promoter complex. The yellow and red diamond indicate the start and end state for the FPT respectively. The figure is adapted from (39).

##### Step 2

While the dwell-time distributions are single exponential for all conditions measured, we learn from the temperature dependency that the system has at least two states. Since *k*_open_ is zero in the limit of zero RNAP concentration, there is no exit rate from the starting state. We therefore conclude that the minimal model describing the transition is an equilibrated two-state system containing only an exit pathway from the intermediate state.

The model consists of an unbound promoter state (state 0) and an RNAP-bound closed-complex state (state 1) that interconvert in rapid equilibrium with forward and backward rate 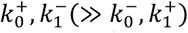. From the RNAP-bound state, the complex transitions irreversibly to the open complex with rate 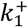, while this is not possible from the unbound state 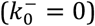.

i. The linear RNAP concentration [RNAP] dependence is on the binding rate 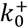 as in **Equation 4**. Using R and P to denote RNAP and promoter, respectively, and RP_c_ to denote the RNAP-promoter closed complex, the effective transition rate for open-complex formation k_open_ is given in terms of [RNAP] by

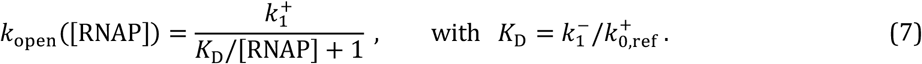

**Equation 7** accurately captures the dependence of *k*_open_ on RNAP concentration obtained from the dwell-time distributions of the CS-to-OS transitions (**Figure 4A**).
ii. The bimodal dependence of the effective CS-to-OS transition rate on temperature is a characteristic signature of an underlying two-state system (**Figure 4B**), despite the single-exponential dwell-time distributions. To quantitatively describe this behaviour, we rewrote the expression for *k*_open_, introducing the temperature dependence of each microscopic rate in terms of its activation energy *E*_*i*_

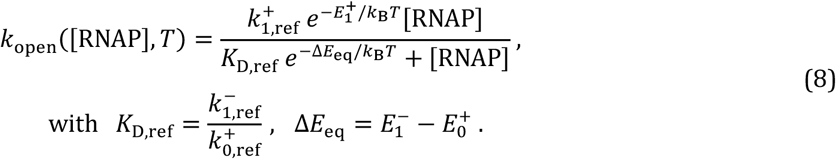

**Equation 8** captures both the observed RNAP concentration and temperature dependence (**Figure 4AB**) (39).

##### Step 3

In general, for a two-state system with a single exit pathway that exhibits a bimodality as a function of temperature fitted with **Equation 8**, 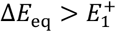 must hold. This was the case for the fitted parameters and also the values for *K*_*D*,ref_ and 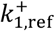 were positively valued (39), confirming that the fit was successful.

Since open-complex formation occurs exclusively from the RNAP-bound state (62), the two equilibrated states in this model correspond to the unbound promoter state and the RNAP-bound closed-complex state and there is only open-complex formation from the closed-complex state.

This interpretation is supported by previous single-molecule studies of RNAP-promoter open-complex formation, which reported that the open-complex formation step is slow compared with the binding and unbinding dynamics of RNAP at the promoter (47, 63, 64).

### OS-to-CS transitions: Kinetic model for the closing of the RNAP-promoter open complex

We now derive the kinetic model describing the transitions from the open state (OS) to the closed state (CS) of the RNAP-promoter complex using the data-oriented model selection procedure (including Step 1c) based on the shifts in the dwell-time distributions with RNAP concentration.

#### Step 1a

The dwell-time distributions for the open-complex closing transitions (OS-to-CS) exhibited two distinct shoulders under all measured conditions, as in **Figure 1H** (39). These distributions were accurately described by double exponential pdf’s with two characteristic timescales, indicating that the underlying kinetic scheme contains at least two non-equilibrated effective states.

#### Step 1b

No significant dependence of the double exponential fit parameters *k*_c1_, *k*_c2_, and *p*_c2_ on RNAP concentration was observed (**Figure 4DE**). This confirms that the measured OS-to-CS transitions originate from single RNAP-promoter open complexes and that the observed kinetics can be described by a two-state system. The presence of two shoulders in the distributions restricts the possible models to a subset of non-equilibrated two-state systems, in which the exit rate from the initial state (state 0) exceeds the exit rate from the intermediate state (state 1) 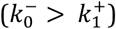 (**Figure 2B**).

#### Step 1c

Since the non-equilibrated two-state system could still have four microscopic rates, while the fits to the distributions are defined by only three, there is no direct mapping of the parameters without a rate dependency. Therefore we determined the minimal kinetic model by combining the data with knowledge from the literature.

Previous studies have shown that RNAP-promoter open-complex formation involves multiple intermediates with distinct stabilities (62). Based on this knowledge, we identified states 0 and 1 in the OS-to-CS transitions as an unstable and more stable RNAP-promoter open-complex, respectively. We considered that the closing from the stable open-complex state is negligible compared to the closing from the unstable open-complex state, so we set the corresponding rate to zero 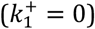.

#### Step 2

The minimal model describing the observed data is a non-equilibrated two-state system containing a single exit pathway from the initial state (state 0) and no dependence on RNAP concentration. Within this framework, the experimentally determined double exponential fit parameters *k*_c1_, *k*_c2_, and *p*_c1_ can be related to the microscopic rates 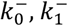 and 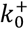 through the following conversion relations (**Methods**)

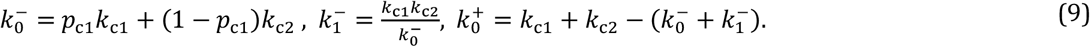

The microscopic rates extracted from the double exponential fit parameters using **Equation 9** also showed no significant dependence on RNAP concentration (**Figure 4DE**), demonstrating that the proposed kinetic model accurately captures the observed dynamics. Together, these results confirm that the two-state model shown in **Figure 4F** provides a complete description of the OS-to-CS transitions (39).

#### Step 3

The microscopic rates extracted from the double exponential fit parameters with **Equation 9** were all positively valued, confirming the fit was successful (39).

Previous studies have shown that RNAP-promoter open-complex formation involves multiple intermediates with distinct stabilities (62). Based on this knowledge, we interpret the OS-to-CS transition as originating from an unstable intermediate open-complex state, denoted RP_I_. From this state, the complex can either close directly and dissociate or undergo repeated transitions to a more stable open-complex state, RP_O_, before eventually returning to the unstable intermediate and dissociating (**Figure 4F**).

This interpretation is supported by another single-molecule study of RNAP-DNA open-complex formation using FRET, which identified an intermediate partially opened state with relatively low stability, whereas the fully opened complex was found to be significantly more stable (47). In that study, direct closing from the stable open state was not observed. Instead, transitions first occurred back to the intermediate state, suggesting that complete closing proceeds through this less stable intermediate. Furthermore, positive supercoiling of DNA (**Figure 1A**) was shown to reduce the stability of the open complex, resulting in an increased overall closing rate (64), in agreement with our observations.

### A course-grained dwell-time analysis for continuously varying time-traces

We now turn to the analysis of dwell-times obtained from continuously varying single-molecule trajectories, such as those recorded for template elongating polymerases and helicases with magnetic tweezers (**Figure 1CD**). In experimental techniques such as magnetic tweezers, the spatiotemporal resolution is often insufficient to resolve the individual steps associated with each nucleotide incorporation cycle or base pair unwinding event. Nevertheless, mechanochemical information about these molecular motors can still be extracted through coarse-grained dwell-time analysis, in which dwell-times are determined over a extended spatial window corresponding to a fixed distance (**Figure 1FI**). In general, deriving the first-passage-time (FPT) distribution analytically for a process involving many stochastic steps quickly becomes very complicated (**Methods**). However, in the special case where the molecular motor proceeds through a sequence of steps, we can often calculate the FPT distribution to complete *N* forward steps, e.g. representing the incorporation of *N* nucleotides or the unwinding of *N* base pairs. Here we assume that we can disregard sequence dependency, which can be supported by overlaying the time-traces obtained for a single experimental condition and confirming that there are no clear sequence dependent trends.

The minimal stochastic model of stepping through a window of size *N* is that the motor performs *N* steps with a constant characteristic rate *k*_f_, or equivalently with characteristic timescale *τ* = 1/*k*_f_ (**Figure 5B**). The FPT distribution for this model is a gamma distribution

**Figure 5:**
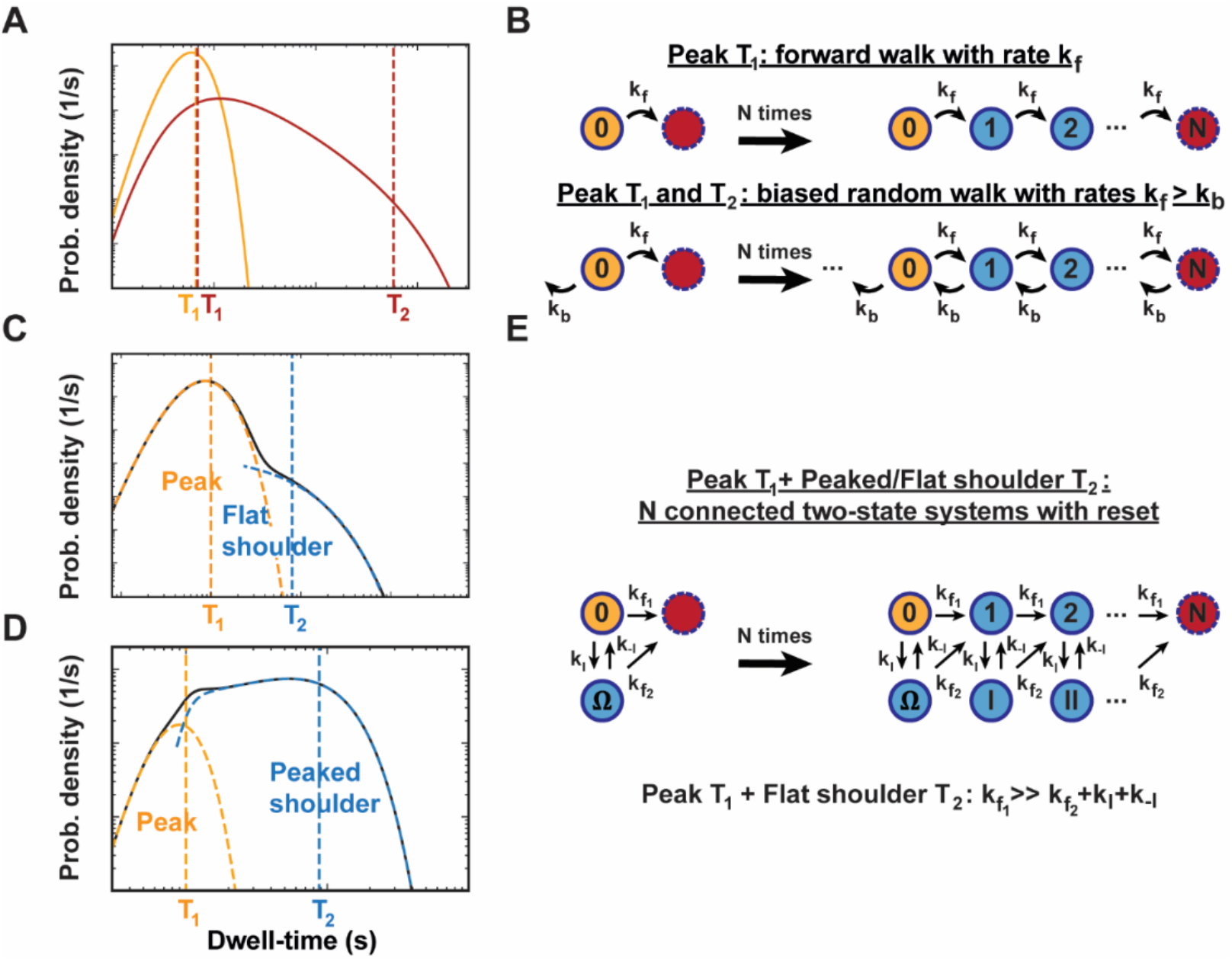
The dwell-time distributions for single- and two-state systems translated to non-overlapping windows of *N* steps. **(A)** In the simplest case, the dwell-time distribution on the window of *N* steps is well-described by a peak with a single characteristic timescale *T*_2_ (yellow curve) and well-fitted with a gamma distribution. Alternatively, the dwell-time distribution has a different characteristic timescale marking the rise *T*_1_ and fall *T*_2_ (red curve). **(B)** The minimal model for a peaked distribution with a single timescale *T*_1_ is an *N* times connected single-state model with forward rate *k*_f_ per step (top). For a peaked distribution in which the rise and fall are characterized by different timescales *T*_1_ and *T*_2_, the minimal model is an *N* times forward-connected single-state model with forward rate *k*_f_ and backward rate *k*_b_(< *k*_f_) on each position, referred to as a biased random walk (bottom). This model also contains an infinite chain of states in the backward direction. **(C)** Dwell-time distribution with a “peak” at short timescales defined by timescale *T*_1_ (yellow dashed curve) and a “flat shoulder” characterized by long timescale *T*_2_ (blue dashed curve). **(D)** Dwell-time distribution with a “peak” for short timescales defined by timescale *T*_2_ (yellow dashed curve) and a “peaked shoulder” with long timescale *T*_2_ (blue dashed curve). **(E)** A dwell-time distribution with two distinct features—a peak with short timescale *T*_1_ and a flat or peaked shoulder with long timescale *T*,(C, D)—can be described by an *N* times forward-connected two-state model that resets to the main state *i* ∈ {0,1,2, . . .} after each step. A peak with short timescale and a long timescale flat shoulder (C) arise when the forward rate from state 0 is larger than the sum of the remaining rates (*k*_f1_ ≫ *k*_f2_ + *k*_I_ + *k*_−I_).

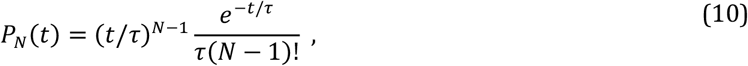

which displays a narrow peak (**Figure 5A**) around the characteristic timescale *T* = *Nτ* with standard deviation 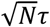 (**Methods**). The short time behavior of this distribution is ∼ *t*^*N*−1^ for *t* < *τ* and for *t* > *τ* the distribution decays as ∼*e*^−*t*/ *τ*^, so this distribution is controlled by a single timescale *τ* and the window size *N*, for which the product is the characteristic timescale *T* = *Nτ*.

Any observed peaked distribution that has two inherent characteristic timescales (**Figure 5A**) indicates that a more complex kinetic mechanism underlies the motor dynamics. A minimal model with identical steps yielding such a distribution type is the forward-biased random walk with a single forward rate *k*_f_ and single backward rate *k*_b_ on each step (**Figure 5B, Methods**). This model has the first-passage time distribution (65)

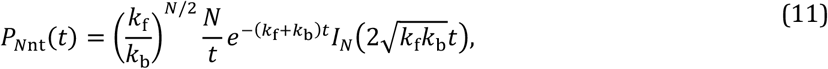

where *I*_*N*_(⋅) denotes the modified Bessel function of the first kind and the condition *k*_f_ > *k*_b_ has to hold for this distribution. The rise of this distribution is characterized by ∼ *t*^*N*^ cut-off at 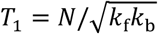 and a decay on long timescales 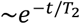 characterized by 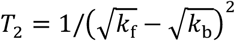, while there is a power law decay ∼ *t*^−3/2^ independent of the rates in the intermediate regime *T*_1_ > *t* > *T*_2_ (66). Note that since there is a backward rate from each state, the chain of states is infinitely long in the negative direction, on which paths can in principle run off. The condition *k*_f_ > *k*_b_ ensures that the paths will always come back to the starting state and eventually make *N* steps forward, and the long paths are characterized by timescale *T*_2_ (65, 66).

Next, we consider the case there are two independent features in the FPT distribution, i.e. a “peak” on short timescales, defined by a gamma distribution with characteristic timescale *T*_1_ and combined with a “flat shoulder” or a “peaked shoulder” with long characteristic timescale *T*_2_ (**Figure 5CD**). The minimal description for these distribution types in our framework is an *N* times connected two-state system with the same rates for each step and a reset to the main state (**Figure 5E**). Here the reset provides that the kinetics of the motor are the same on each step. The “flat shoulder” is obtained only when *k*_f1_ ≫ *k*_f2_ + *k*_I_ + *k*_−I_ (**Methods**), while there is a “peaked shoulder” in the distributions for all other regimes of the microscopic rates.

The FPT distribution for this model could in principle be numerically solved using the Master Equation approach (**Methods**), but this is computationally costly. Instead, we can determine the features in the dwell-time distribution and construct an interpolated fit-function that captures the dominant features and their connection to the microscopic rates.

When the characteristic features are clearly distinguished in the dwell-time distribution *T*_2_ ≫ *T*_1_ (**Figure 5CD**), we can assume separation of timescales for the underlying two-state system for single nucleotide steps (*τ*_2_ ≫ *τ*_1_), where *τ*_1_(= 1/*k*_c1_) and *τ*_2_(= 1/*k*_c2_) are the characteristic timescales from the double exponential distribution for a single step in the model (**Equation 1**). As explained in the **Methods**, in this case we can split the expression for the FPT distribution in the term for *N* steps with timescale *τ*_1_ and the term for the combinations of paths with at least one step with timescale *τ*_2_, disregarding the contributions of timescale *τ*_1_ in the second term

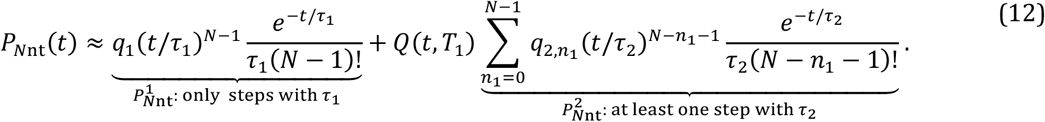

The first term 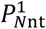 is a gamma distribution with average timescale *T*_1_ = *Nτ*_1_ and weight 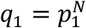. The second term 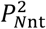 is the sum of gamma distributions with index *n*_1_, average timescale 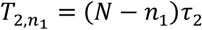 and binomial weight 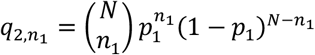, which yields either a flat or peaked shoulder. We also consider here that 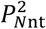 is an approximation that only works for long timescales, since the distribution rises as ∼ *t*^*N*−1^ on short timescales up to the first peak with characteristic timescale *T*_1_ (**Methods**). Therefore, we added a cut-off factor *Q*(*t, T*_1_) at time *T*_1_ to the term 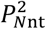 ensuring that the short time behaviour follows the initial rise

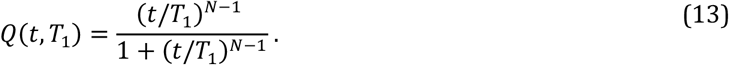

In case the shoulder on long timescales is “flat” and well-fitted by a single exponential distribution (**Figure 5C**), the term in 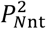 representing the paths with a single step at timescale *τ*_2_ dominates and therefore 0 < *p*_2_ ≪ *p*_1_. With separation of timescales *τ*_2_ ≫ *τ*_1_, we also obtain a condition on the microscopic rates for the “flat” shoulder: *k*_f1_ ≫ *k*_f2_ + *k*_I_ + *k*_−I_ (**Methods**).

Since all the gamma distributions in the term 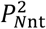 show the same decay 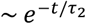, they form a single flat shoulder on long timescales in this limit, which is well-described by a single exponential approximation defined by the overall weight of the shoulder *q*_2_ and average timescale *T*_2_ for all combinations with at least one step at timescale *τ*_2_ in terms of single step parameters (**Methods**)

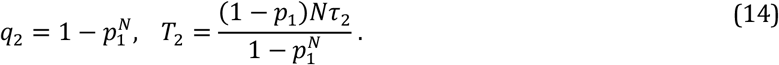

With the single exponential approximation of 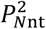 and the cut-off factor for short timescales at *T*_1_ (**Equation 13**), we arrive at a fit-function that captures the peak and shoulder in the dwell-time distribution

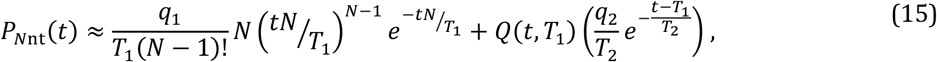

with *T*_1_ = *Nτ*_1_ and 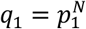 while the expressions for *T*_2_ and *q*_2_ are given in **Equation 14**.

While **Equation 12** and **15** are normalized over an infinite time window, the window is limited in reality. Since we typically trust events only longer than some multiple of our time resolution, we employ a lower time cut-off *t*_min_. And when maximum likelihood estimation (MLE) is used for fitting, it is important to make sure that the total probability of the dwell-times adds up to 1, so we strictly enforce it by

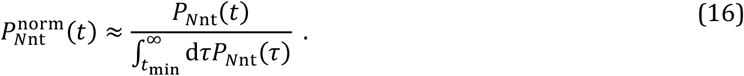

With this normalization, the approximated FPT distribution **Equation 12** with **Equation 13** can be used as a fit-function to dwell-time distributions consisting of a peak at short timescales and a flat/peaked shoulder on long timescales or **Equation 15** with **Equation 13 and 14** specifically for dwell-time distributions consisting of a peak at short timescales and a flat shoulder on long timescales. From these fits we obtain the timescales *τ*_*i*_ and probabilities *p*_*i*_ in the underlying two-state system for single steps, from which we can check the separation of timescales *τ*_2_ ≫ *τ*_1_ and in case of a flat shoulder the separation of probabilities *p*_2_ ≪ *p*_1_. The values of all microscopic rates of the minimal kinetic model (**Figure 5E**) can be obtained by performing a concentration or force sweep and fits to the dwell-time distributions to extract the values for *τ*_*i*_ and *p*_*i*_ at the measured conditions (**Methods**). With this information, the underlying limiting case of the two-state system can be determined for single steps by the motor (**Figure 3**).

In the coarse-grained dwell-time analysis of molecular motor traces, we often observe a peak at short timescales, fitted with a gamma distribution, in combination with more than one flat shoulder on long timescales that can be fitted with exponential distributions (**Figure 1I, Methods**) (18, 29, 57, 61). In such cases, this coarse-grained modelling framework can be extended to two or more alternative states per step as described in studies on polymerase activity (18, 29, 57, 61). Additionally, **Equation 12, 14 and 15** could be similarly expanded for fitting more flat/peaked shoulders with separated timescales (**Methods**).

### Graphical user interfaces (GUIs) to perform dwell-time analysis and MLE fitting

To help with applying our approach for data processing and analysis, we developed graphical user interfaces (GUIs) that enable both data sorting and processing to generate dwell-time distributions and MLE fitting for complex kinetic models and statistical error estimation.

For the processing of single-molecule time-traces, we distinguish two types: discrete and continuous time-traces. For the discrete time-traces on the opening and closing of the RNAP-promoter complex illustrated in **Figure 1E**, Bera et al. (2022) used a change point detection method to detect the step transitions in the trace from the python package Ruptures (67). The change point analysis was implemented in a set of GUIs, in which reference bead trace subtraction, trace part selection, noise Viltering, change point detection and validation and dwell-time extraction can be performed (https://gitlab.com/DulinlabVU/change_point_analysis). Next to change point analysis, HMM analysis is also implemented in this program.

For the coarse-grained dwell-time analysis of continuous time-traces from magnetic tweezers assays on the nucleotide addition cycle of polymerases and nucleic acid unwinding activity of helicases (18, 29), we have another GUI program. The program includes the tools for reference bead trace subtraction, trace part selection, noise filtering, dwell-time detection and validation (https://gitlab.com/DulinlabVU/magnetic_tweezer_trace_analysis).

To perform MLE fitting of the probability density function (pdf) curves on the dwell-times distributions, we provide a GUI program that includes the pdf curves of all the distribution types discussed in this manuscript (https://gitlab.com/DulinlabVU/mle_fitting_dwell_time_distributions). In this program, short and long time cut-offs with renormalization can be applied, the expected parameter ranges can be selected and MLE fitting is performed with basin hopping (68). The statistical error on the fit parameters can be determined by bootstrap fitting, i.e. fitting to resampled sets of the dwell-times.

In case there is not enough information to narrow down a single model with the minimal number of free parameters, fits with the set of remaining models to the dwell-time distributions can still be performed for the measured experimental conditions to select the model that fits best. For this model selection, we recommend weighing the log-likelihood *LL* (goodness-of-fit) with the number of parameters *k* and the number of dwell-times *M*. This can be done by evaluating the Bayesian Information Criterion (BIC) (59)

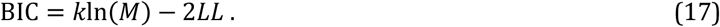

The model that minimizes the BIC is considered the best fit and this value is given in the MLE fitting program. If changes in BIC are not significant, i.e. within one standard error of the mean, the model fit that captures all clear trends while minimizing the number of parameters is preferred. Note that a significant change in BIC for models with similar log-likelihood but different number of parameters can only be observed when a sufficiently large *M* has been collected.

## Discussion

The ability to track individual biomolecules combined with the development of high-throughput single-molecule assays offers the opportunity to capture and reveal the full dynamics of biomolecular reactions. A key aspect of extracting such dynamics is the ability to determine the most accurate model. Such task is generally integrated through multiple rounds of hypothesis formulation, literature search, model fitting, and testing (**Figure 4**). Yet, building a kinetic model has become common practice in single-molecule biophysics and gives valuable insights on the studied reaction (28–31, 54, 57, 69). Here, we propose a systematic approach to model selection based on single-molecule data, starting from the features encountered in dwell-time analysis. If the data at hand contains enough information, data-driven model selection can be performed, requiring only information from the literature to interpret and validate the model. Otherwise, depending on the quality of the data (e.g. spatiotemporal resolution, measurement time, statistics) and limitation on the range of experimental conditions (e.g. only assisting or opposing forces), the data should be supplemented with information from the literature, referred to as data-oriented model selection.

In this manuscript, we described how to extract dwell-time distributions from time trajectories typically obtained in single-molecule experiments. These we divided in two types: the one showing clear discrete steps and the other not, so they are treated as continuous traces (**Figure 1**). For the time-traces with clear steps, we can learn the minimal number of states in the system, the rate dependencies and how the exit rates compare from the type of dwell-time distributions with a single rate dependency on a concentration or force, as illustrated for two-state systems in **Figure 2, 3 and S1-3**.

We discussed that when the steps of a molecular motor are smaller than the experimental resolution, we can still perform coarse-grained dwell-time analysis (**Figure 1FI, Figure 5**). In this analysis, however, we have to assume a constant step size, which may not be the case for the motor of interest. Alternatively, continuous traces could be analyzed by determining changes in the slope of the trace. With such analysis, the pause-free average rate can be obtained (70, 71). To investigate more details in the dynamics, a distribution of the slopes can be built, from which peaks can be distinguished, corresponding to different modes of the motor (17, 71, 72). However, the peaks in the distributions can be hard to distinguish as a result of varying noise levels in the traces and transition modes, so if the step size of the motor is known we recommend performing our coarse-grained dwell-time analysis for more detailed kinetic analysis (**Figure 5**).

For the scope of this manuscript, we covered single- and two-state systems and repeated arrays of them, for which the dwell-time distributions could be analytically solved (**Methods**). Using the Laplace transform method, at least all systems with up to four states and repeated arrays of them with resets could be solved analytically, limited by the order of the polynomials in the transformed distribution (73). In more complex cases, however, the dwell-time distributions can become very hard to solve. Then the FPT distribution can still be calculated numerically using the Master equation approach (**Methods**) (22, 31).

High-throughput single-molecule biophysics promises to give a whole new perspective on complex reaction kinetics in molecular biology. With the improvements of spatiotemporal resolution (14, 74–77), throughput and the development of combined single-molecule techniques with other biochemical assays (11, 13, 78), data with higher order complexity and more intrinsic information will become available soon in this Vield. These developments call for the implementation of systematic and data-driven model selection approaches for higher-order mechanistic models, which could be automated and may be described as a form of ‘interpretable artificial intelligence’.

With the methodology presented, we aim to help single-molecule biophysicists to process and analyze the typical time-traces acquired with these assays when a sufficient number of events has been collected, i.e. >100, enabling dwell-time analysis with high statistics. Our characterization of single- and two-state systems (**Figure 2, 3 and S1-3**) and repeated arrays of them (**Figure 5**) should be viewed as a starting point for systematic kinetic modelling of complex reactions in context of single-molecule traces.

## Data and code availability

This manuscript is supplemented with three data analysis programs in python as discussed in the main text. These are available at https://gitlab.com/DulinlabVU/change_point_analysis, https://gitlab.com/DulinlabVU/magnetic_tweezer_trace_analysis and https://gitlab.com/DulinlabVU/mle_fitting_dwell_time_distributions.

## Acknowledgements

DD was supported by BaSyC – Building a Synthetic Cell” Gravitation grant (024.003.019) of the Netherlands Ministry of Education, Culture and Science (OCW) and the Dutch Organization for Scientific Research (NWO), and NWO funding OCENW.XL21.XL21.115. We thank Thomas Bugea for reviewing the manuscript and Mohammad Sadegh Feiz for reviewing the methods section before submission. We thank for Subhas C. Bera for providing the GUI program for dwell-time analysis on time-traces with discrete time steps. We thank Subhas C. Bera and Mona Seifert for the initial implementations of the GUI program for dwell-time analysis on continuous time-traces and Pauline van Nies for the initial implementation of the GUI program for MLE fitting of dwell-time distributions.

## Author contributions

MD, DD and PA designed the outline of the research. PA drafted the manuscript. MK developed the GUI program for dwell-time analysis of continuous traces, PA developed the GUI program for MLE fitting of dwell-time distributions. PA, MK and MD contributed the theory framework. PA, MK, MD and DD wrote the manuscript.

## Declaration of generative AI and AI-assisted technologies in the manuscript preparation process

During the preparation of this work the authors used ChatGPT in order to improve the structure and language of the manuscript. After using this tool/service, the authors reviewed and edited the content as needed and take(s) full responsibility for the content of the published article.

## Methods

### Laplace transform method to solve first-passage time (FPT) distributions

In the context of single-molecule experiments, dwell-times often represent “the time it takes to complete a reaction for the first time”, also referred to as the first-passage time (FPT) (54). We will now give a convenient method to solve the type of FPT distributions starting from the underlying kinetic scheme. When working with non-equilibrated multi-process models, it is convenient to write the FPT distributions in Laplace space, because convolutions arising when calculating the probability density of the total time to perform sequential steps simply become multiplications (4).

First, consider that the transition-times for a single step of a process from state *A* to state *B* with kinetic rate *k*_*A*→*B*_ follow an exponential decay distribution

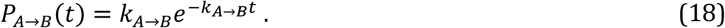

The Laplace transform of this distribution in terms of variable *s* is

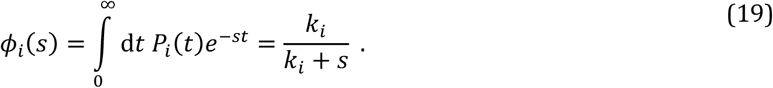

Since we only evaluate real-valued non-negative *s* in this transform 0 < *ϕ*_*i*_(*s*) ≤ 1 holds.

As an example to illustrate the Laplace transform method, we will show how to calculate the FPT distribution for a series of two non-reversible processes. For example, the binding of substrate *S* to an enzyme *E*, followed by the transformation of *S* into product *P* with respective rates *k*_1_ and *k*_2_

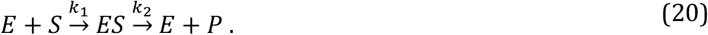

The FPT distribution of the reaction described in **Equation 20** is a convolution of the transition-time distributions for the two individual steps. However, when taking the Laplace transform, the total FPT distribution becomes the product of the two transition-time distributions of the steps and reads

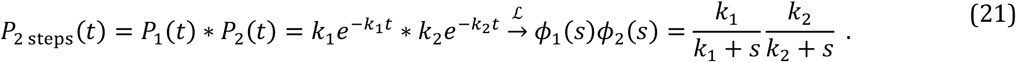

From here, we can use partial fraction decomposition to write the product of fractions as a sum of fractions by introducing probabilistic weights *p*_c1_ and *p*_c2_ = 1 − *p*_c1_

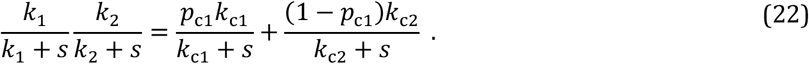

By taking the inverse Laplace transform of the right-hand side, we obtain that the FPT distribution is the sum of two exponential distributions

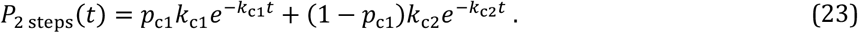

Now we need to find an expression for *p*_c1_ and *p*_c2_ = 1 − *p*_c1_. To this end, we rewrite the right hand side of **Equation 22** as a single fraction

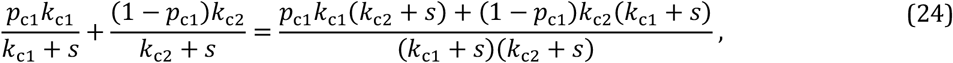

Note that the right-hand side of **Equation 24** can be equalized with the left-hand side of **Equation 22**, when we take for the characteristic rates *k*_c1_ = *k*_1_ and *k*_c2_ = *k*_2_. By comparing the numerators in the left- and right-hand side of **Equation 24**, we obtain from the *s*-term that *p*_c1_*k*_c1_ + (1 − *p*_c1_)*k*_c2_ = 0, from which we solve:

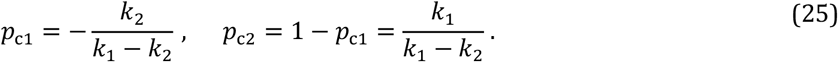

And by substituting the terms in **Equation 25** into **Equation 23**, we obtain the following expression for the FPT distribution

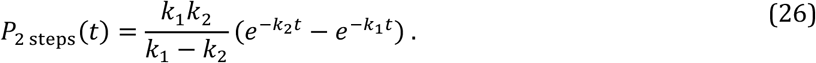

We find that this FPT distribution is peaked (**Figure 2A**), which is generally the case for the difference of two exponential distributions (*p*_c1_ < 0). In case the probabilistic weights for both exponential distributions have positive values (0 < *p*_c1_ < 1), the FPT distribution shows two shoulders on log-log scale (**Figure 2A**). We should, however, keep in mind that **Equation 26** only holds when *k*_1_ ≠ *k*_2_. In the special case that *k*_1_ = *k*_2_, we obtain a gamma distribution of order 2 (**Equation 10** with *N* = 2).

### Derivation of first-passage time (FPT) distributions for non-equilibrated two-state systems

As explained in the main **Results**, all limiting cases of non-equilibrated two-state systems can be distinguished from the types of FPT distributions when we modulate a concentration or a force acting on a single rate in the system (**Figure 3, Figure S1-S2**). In this section, we will derive the relation for the regimes with different distribution types for the complete two-state system and give the set of conversion relations from the two exponential fit parameters to the microscopic rates of the system. Then we discuss the transition points in the probabilistic weight *p*_c1_ and in the characteristic rates of the exponential decays (*k*_c1_ and *k*_c2_) in the two exponential distributions.

### FPT distributions for the complete two-state model starting from state 0

Here we derive the first-passage time (FPT) distribution of the complete two-state model starting from state 0 with an arbitrary number of forward transitions with 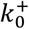 and backward transitions with 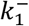 respectively and exit rates from both state 0 and 1 are considered with rate 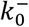 and 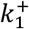 respectively (**Figure 2B**). Considering that there are multiple possible paths from both states, the transition-times show an exponential decay with the sum of rates from the particular state ∑_*j*_ *k*_*j*_ and the transition-time distributions are multiplied by the splitting probability for taking the path 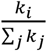, where *k*_*i*_ is the rate for the specific path

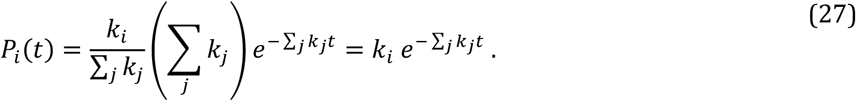

Using **Equation 19**, the particular transition-time distributions for each kinetic step in the two-state system read in Laplace space

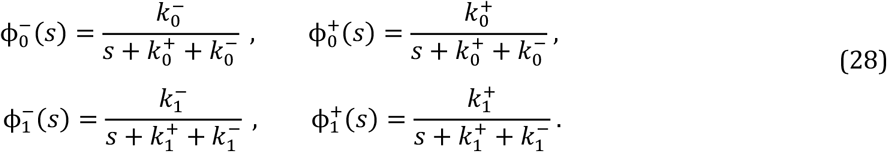

Considering the model starts from state 0, there can either be a direct exit with distribution 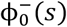, a transition to state 1 and exit from there, yielding the distribution 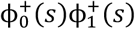, or there is an arbitrary number of forward and backward transitions with distribution 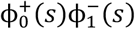 before exit from either state. With the transition-time distributions in **Equation 28**, we obtain the Laplace transform of the FPT distribution

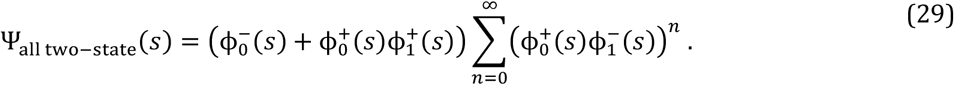

Using that for the geometric sum 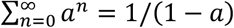 if |*a*| < 1, which is the case for the transition distributions ϕ_−_ and their products (from **Equation 21**), the expression can be simplified to

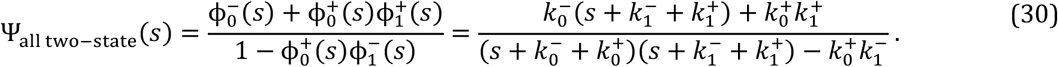

When we expand the denominator on the left hand side of **Equation 30** to the form (*s* + *k*_c1_)(*s* + *k*_c2_), where *k*_c1_ and *k*_c2_ are the characteristic rates, we obtain them by solving the roots of the denominator:

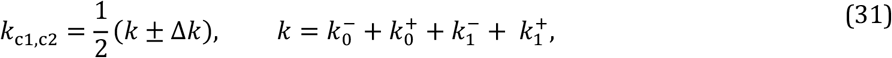

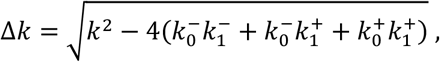

where *k*_c1_ ≥ *k*_c2_.

Using partial fraction decomposition, we obtain that the FPT distribution consists of two exponential distributions with rates *k*_c1_ and *k*_c2_ and probabilistic weights *p*_c1_ and 1 − *p*_c1_

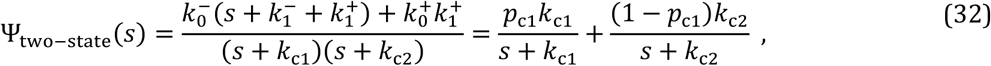

which holds for 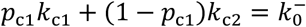 and is solved for

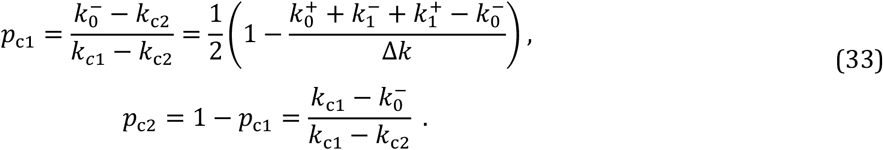

To obtain conversion relations for the two exponential fit parameters (*k*_c1_, *k*_c2_ and *p*_c1_) to the microscopic rates of the complete two-state model, the right-hand side of **Equation 32** is rewritten as a single fraction, as in **Equation 24**. By comparing the numerator of the resulting expression with the numerator on the left-hand side of **Equation 32**, we obtain the following set of relations:

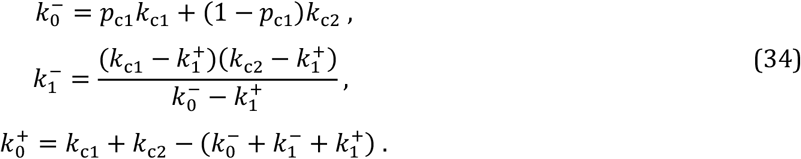

A unique expression for 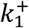 in terms of the fit parameters *k*_c1_, *k*_c2_, *p*_c1_ and microscopic rates 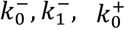 can however not be found, so this is not a full set of conversion relations.

### Obtaining a full set of conversion relations for the microscopic rates to the fitted parameters

However, in the case that *C* (or *F*) is acting on 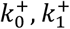 can be found as the single exponential rate in the limit of very high *C* (or *F*) (**Figure 3C**), giving a full set of conversion relations.

When the *C* dependency is on 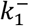, the distribution at *C* = 0 will present two characteristic timescales (i.e. two shoulders or a peak) of the same type as the distributions at intermediate *C*^∗^ and is well-fitted with a two exponential distribution with parameters 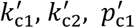, which give the following relations for the internal rates (using 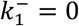)

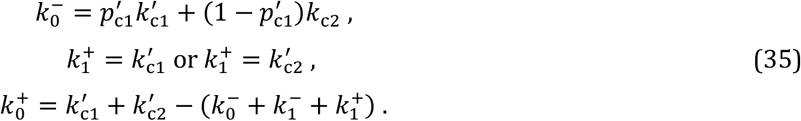

From the second relation 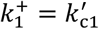 or 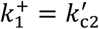, depending on whether 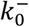 is greater than 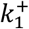 or the reverse.

### Transition points in the two exponential distributions

We define a transition point as the condition on the parameter at which the FPT distribution is single exponential and the distribution changes shape for slightly different parameters.

### Transition points in the probabilistic weight of the first exponential p_c1_

The probabilistic weight *p*_c1_ shows a transition point at *p*_c1_ = 0, where the FPT distribution reduces to a single exponential. Namely for 0 < *p*_c1_ < 1, the FPT distribution consists of two exponential components with positive amplitudes, resulting in a double-exponential decay. In contrast, for *p*_c1_ < 0, the distribution becomes peaked due to the negative contribution of one exponential component.

From **Equation 33**, we find that the condition *p*_c1_ = 0 is satisfied when 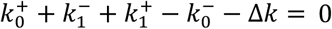. This condition simplifies to either 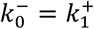 or 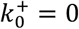. The latter case corresponds to the absence of forward transitions from state 0 and therefore results only in single-exponential FPT distributions, rather than a transition between the two regimes. Consequently, the transition point *p*_c1_ = 0 is reached for 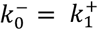.

We obtain *p*_c1_ > 0 when 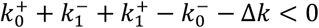, which reduces to 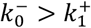, whereas *p*_c1_ < 0 follows from 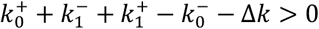 corresponding to 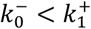.

In the regime 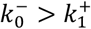 both exponential contributions have positive weights, yielding a double-exponential FPT distribution. At the boundary 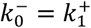 the distribution becomes single exponential, whereas for 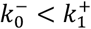 the first probabilistic weight *p*_c1_ is negative, producing a peaked FPT distribution.

### Transition points in the two exponential rates

In principle, there is a transition point at *k*_c1_ = *k*_c2_ for all non-equilibrated two-state systems. However, considering that the transition point can be rewritten to Δ*k* = 0, which is satisfied when 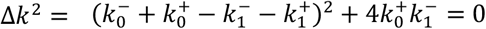 (from **Equation 31**) and reduces to 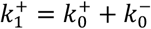, it can only be reached if 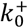 or 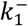 is zero. However, 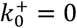 is not a transition point, as discussed above.

The transition point at 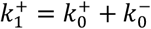 is reached only if 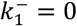, we modulate a concentration or force acting on the nonzero rate 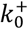 and there are more exits from state 1 than from state 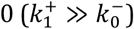. The transition point is still reached if, additionally, 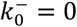 holds, but then at 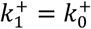.

### Master equation approach for solving FPT distributions

While the Laplace-transform method is convenient for analytical derivations in low-dimensional systems, the Master equation offers a more general and numerically tractable framework for computing first-passage time distributions in complex kinetic networks.

Considering that the changes in fractional occupancy *P*_*i*_ of states *i* ∈ {1,2} at timepoint *t* for a single decay process with rate *k* from state 1 to state 2 reads

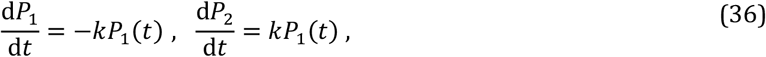

where *kP*_1_(*t*) represents the “probability flux” from state 1 to state 2 at timepoint *t*.

The FPT distribution of the decay is obtained as the time derivative of the fractional occupancy of state 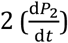. For this, we should first solve the fractional occupancy of state 1 *P*_1_ (*t*) from the first expression in **Equation 36**, which can then be substituted in the second expression

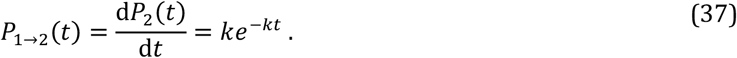

In the case we have multiple processes connecting a set of *N* states, we obtain a set of differential equations, describing the change in fractional occupancy of each state at timepoint *t*. To condense the set of equations, we can write the fractional occupancies of the states at timepoint *t* as a column vector 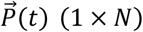. Multiplication of 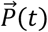 with the transition rate matrix *M* (*N* × *N*) containing all rates from every state gives the changes in the fractional occupancies of every state at timepoint *t*, which is generally referred to as the Master equation

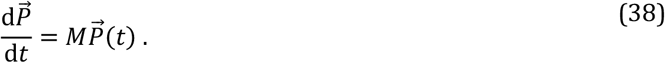

In the transition rate matrix *M*, the off-diagonal elements contain transition rates between states and the diagonal elements ensure probability conservation, as exemplified for the complete two-state system in **Equation 41**. The formal solution of the Master equation is given by

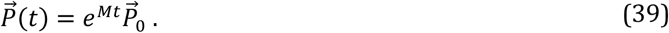

The first-passage time distribution to a designated target state is obtained from the probability flux into that state, i.e. as the sum of incoming transition rates weighted by the occupancies of the originating states, given by the corresponding element of 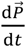 from **Equation 38**.

While the Master equation framework can in principle be applied to arbitrary kinetic networks, numerically solving the resulting system of differential equations is more computationally intensive than calculating the analytic solution, which can be an issue when performing model fits to data. The Laplace transform approach described above is preferred for deriving analytical expressions in systems with a small number of states. The distributions for models with at least up to four states can be solved analytically, limited by the order of the polynomials in the transformed distribution (73). In contrast, the Master equation formulation is particularly well-suited for numerical evaluation in matrix form, and becomes the method of choice when analytical solutions are intractable (30, 31). The Master equation approach is also preferred when there is at least one internal equilibrium in the system.

### Derivation of first-passage time (FPT) distributions for equilibrated two-state systems

#### System of equations for complete two-state system

The complete two-state system can in principle also be solved using the master equation approach, which is the preferred method when there is an internal rapid equilibrium. We start by writing the system of equations for the changes in fractional occupancies with time for each state in the complete two-state system including the exit states (state -1, 0, 1 and 2), which are given by the sums of probability fluxes as exemplified in **Equation 36**

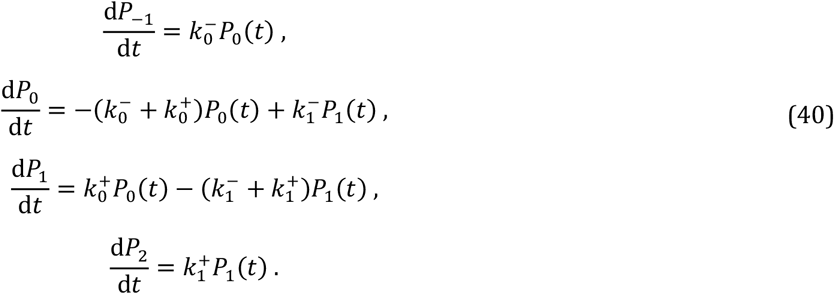

When we write the complete system of equation in matrix form it could in principle be solved using linear algebra as explained in Ref. (1)

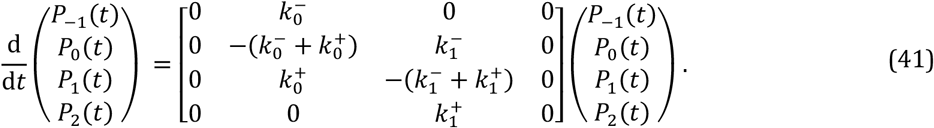

We choose to derive the distribution for the FPTs starting from state 0 to states -1 and 2 combined using the Laplace transform method as explained in the section “**Laplace transform method to solve first-passage time (FPT) distributions**”. In case of a rapid equilibrium between state 0 and 1, the system of equations reduces to a simple set, which can be solved without any use of matrices or the Laplace transform, as shown in the following section.

#### FPT distribution for two-state models with rapid equilibrium between the states

When we consider the rates for transitioning between the states are very large compared to the exit rates, we can assume that the transitioning is equilibrated before the exit from one of the states

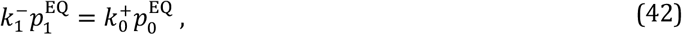

which is referred to as the detailed balance and can be rewritten to 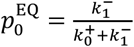 using that 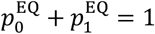.

With the detailed balance, state 0 and 1 can be treated as a single state with regard to the exit of the system, giving rise to a single shoulder in the FPT distribution with characteristic rate 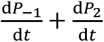. The characteristic rate is expressed as

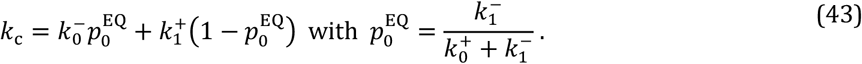

Which simplifies to

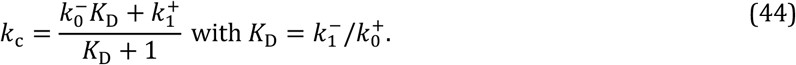

When there is no exit rate from state 1 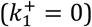 (**Figure 2B**), the expression simplifies to

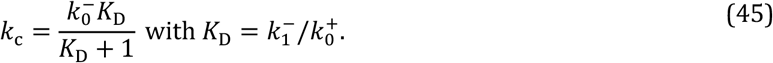

#### Distinct trends for equilibrated two-state models when modulating the microscopic rates Concentration dependency acting on 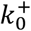

Consider a two-state system in which the forward internal rate depends on a controlled concentration *C*, such that 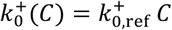 (**Equation 4, Figure 3C**). If only single-exponential FPT distributions are observed over the entire range of measured concentrations, the underlying kinetics can be described either by a single-state model or by a two-state system that is in rapid equilibrium (**Figure 2B**). In the following, we consider the latter case, where the internal transitions equilibrate rapidly compared to the timescale of the measured process.

For the complete two-state system with a rapid equilibrium the expression for the characteristic rate with concentration *C* acting on 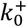 reads

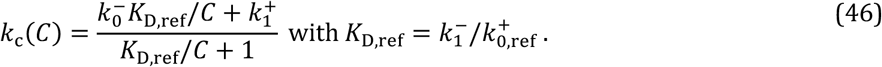

This expression describes a monotonic transition between the limiting rates 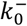 and 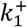. At zero concentration, the characteristic rate approaches 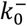, whereas at high concentration it approaches 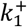. Consequently, the concentration dependence results in an increase or decrease of the characteristic rate depending on whether 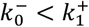 or 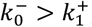, respectively.

#### Concentration dependency acting on 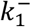

Alternatively, when we modulate a concentration *C* acting on the backward internal rate 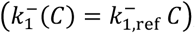, we obtain for the characteristic rate of the equilibrated two-state systems

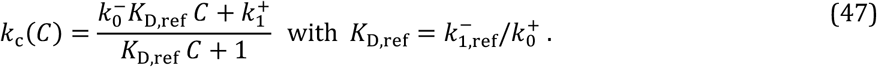

In this case, the concentration dependence results in an increase or decrease of the characteristic rate depending on whether 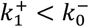 or 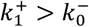, respectively.

#### Force dependency acting on 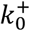 or 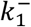

Alternatively, equilibrated two-state models can be distinguished by applying an external force *F* that affects the internal forward transition rate 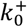 or the backward rate 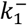, taking the effective *F* in the forward (+) direction (**Figure S2**). For an Arrhenius-type force dependence *F* on 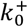 or 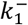(**Equation 5**), the force-dependence of the characteristic rate are given by

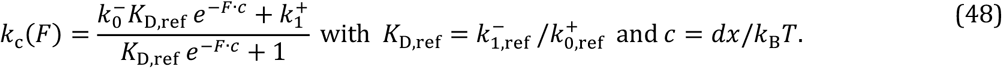

Due to the exponential force dependences on the rapidly equilibrated rates *K*_*D*_, the characteristic rate displays a sigmoidal force dependence, transitioning from 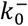 at large opposing forces to 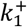 at large assisting forces (**Figure S2A**). If either 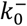 or 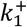 is zero, the corresponding asymptotic limit of the sigmoidal curve approaches zero.

#### Distinct trends in the characteristic rate with temperature for two-state systems in rapid equilibrium

In this section, we investigate the mechanism underlying the non-monotonic temperature dependence of the RNAP–promoter opening rate *k*_open_, which exhibits an increase followed by a decrease with temperature (**Figure 4B**). The observation of a single shoulder in the FPT distributions, a zero opening rate *k*_open_ at zero RNAP concentration, and a bimodal temperature dependence indicates that *k*_open_ is governed by an equilibrated two-state model with a single exit pathway (**Figure 4C**). We therefore analyze the temperature dependence of the characteristic rate for this model, considering the limiting case 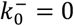 and then we infer the limits on the parameters in the case 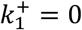.

#### Trends in characteristic rate with temperature for the equilibrated model with 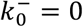

The corresponding expression for the temperature *T* dependent characteristic rate can be written as:

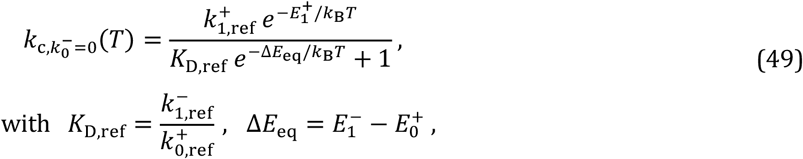

where *K*_*D*,ref_ is the equilibrium constant for the transitioning between state 0 and 1 in the limit of very high temperature and *ΔE*_eq_ is the difference in activation energy for the backward to the forward transitions between state 0 and 1.

To Vind the maximum/minimum in *k*_c_(*T*), we first take the derivative to *T*

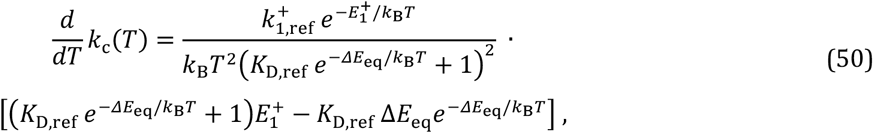

which can be simplified to

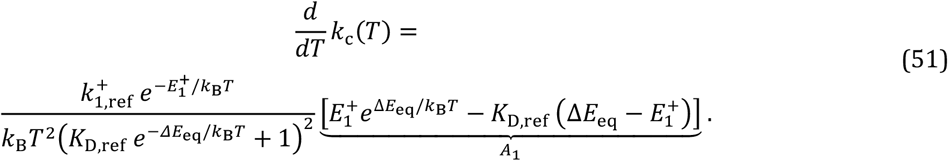

At the maxima and minima of *k*_c_ (*T*), we have 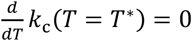. Considering that the first term in **Equation 51** is nonzero for finite *T*, we obtain that the second term *A*_1_ = 0

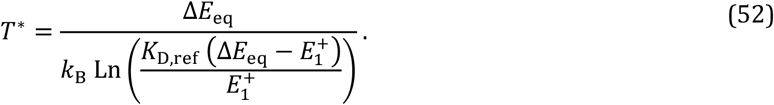

The maximum/minimum is obtained for Δ*E*_eq_ > 0 and 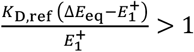, which simplifies to 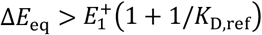 and requires Δ*E*_eq_ > 0. Considering that 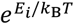 is monotonically decreasing with *T* for *E*_*i*_ > 0, for *T* < *T*^∗^ the second term in **Equation 51** is positively valued (*A*_1_ > 0), while for *T* > *T*^∗^ it is negatively valued (*A*_1_ < 0), meaning that *T*^∗^ is a maximum in *k*_c_(*T*) when Δ*E*_eq_ > 0 (**Figure S3**).

#### Trends in characteristic rate with temperature for the equilibrated model with 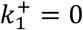

When 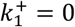, the expression for the temperature dependency of the characteristic rate simplifies to

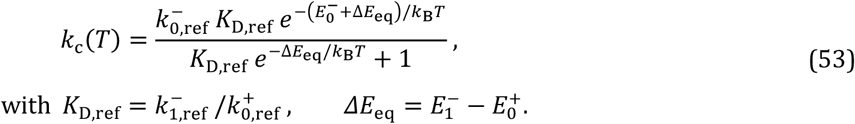

Consider that **Equation 53** can be inferred from **Equation 49** by interchanging state 0 and state 1. Therefore we directly obtain from **Equation 52** that the maximum in the characteristic rate *k*_c_(*T*) with *T* is at

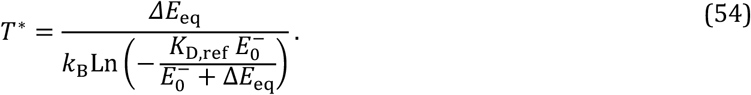

Using the same arguments as for **Equation 52** with state 0 and 1 interchanged, the maximum at *T*^∗^ is reached when Δ*E*_eq_ < 0 and 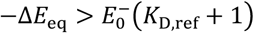 (**Figure S3**).

#### Single exponential approximation of FPT distributions

In some cases kinetic models become too complex to solve analytically, for example when the polynomials in the numerator and denominator of the Laplace transformed FPT distribution Ψ(*s*) becomes too high-order to obtain its roots explicitly (at least up to fourth order polynomials can be generally solved (73)). Then the FPT distribution can instead be approximated by a single exponential characterized by the weight *A* and characteristic timescale *T*_c_. These parameters can be obtained directly from the Laplace transform of the FPT distribution:

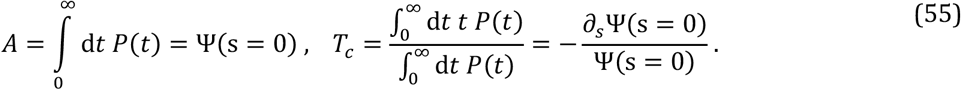

This approximation is useful when the full expression of the FPT distribution is not required and the dominant timescale is sufficient to describe the process. Examples include estimating the characteristic rate *k*_c_, or describing dwell-time distributions (or parts thereof) that exhibit an approximately exponential shoulder on a log-log scale, such as the nucleotide addition cycle of viral RNA polymerases (**Figure 1I**).

To demonstrate the calculation, consider a sequential two-step process with rates *k*_2_ and *k*, (see **Methods** section “**Laplace transform method to solve first-passage time (FPT) distributions**”). For this example we obtain from **Equation 23**

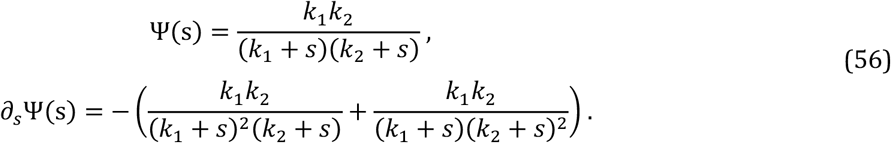

Thus corresponding weight and characteristic timescale are given by:

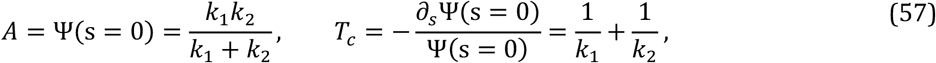

By inverting the characteristic timescale, the characteristic rate for two processes in series is

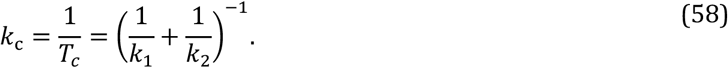

#### The FPT distribution for N steps with the same rate

The minimal description for motor activity on a window of *N* steps forward is that it performs those *N* steps with a constant rate *k*_f_, or equivalently with characteristic timescale *τ* = 1/*k*_f_ (**Figure 5B**). To derive the corresponding first-passage-time (FPT) distribution, we make again use of the fact that convolutions in the time domain become products in Laplace space. The dwell-time distribution for a single step is an exponential decay with timescale *τ* and its Laplace transform is given by 1/(1 + *τs*). The FPT distribution for completing *N* consecutive steps is therefore calculated in Laplace space as

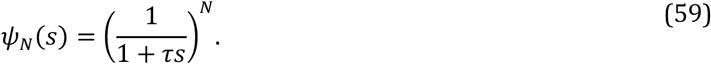

Applying the inverse Laplace transform yields a peaked FPT distribution corresponding to a gamma distribution with shape parameter *N* and timescale per step *τ* (**Figure 5B**) (54)

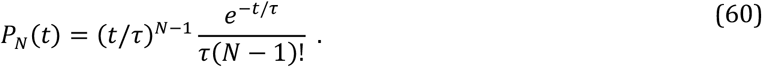

This distribution is defined by the average timescale *T* = *Nτ* and standard deviation 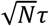.

#### Derivation of the FPT distribution for N passes through identical two-state systems

To derive the distribution for *N* passes through identical two-state systems, first consider that the distribution for a single step is two exponential (**Equation 1**). Then the Laplace transform of the distribution over a window of *N* consecutive steps is

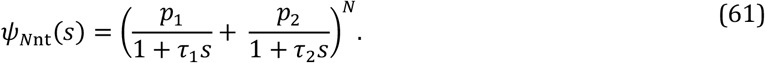

When assuming separated timescales *τ*_2_ ≫ *τ*_1_ we can separate out the terms for *N* steps with only timescale *τ*_1_ and the combinations of paths with at least one step with timescale *τ*_2_

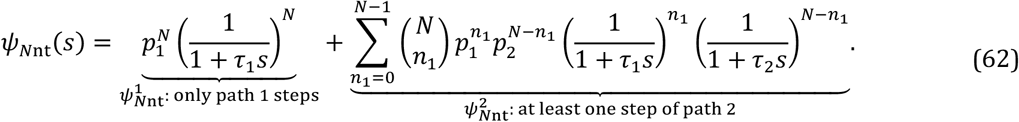

The inverse Laplace transform of the first term 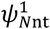 gives a gamma distribution with average timescale *T*_1_ = *Nτ*_1_ and weight 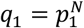. Upon the inverse Laplace transform of the second term, however, we obtain a sum of convolutions of two gamma distributions with order of *n*_1_ and *N* − *n*_1_ times the weights 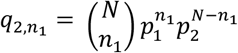, which is compute-time consuming and not convenient to calculate during a fitting procedure. Nevertheless, when there are two distinguishable features in the dwell-time distribution, i.e. a peak at short timescales and a flat/peaked shoulder at long timescales (**Figure 5CD**), we can assume a separation of timescales *τ*_2_ ≫ *τ*_1_. Then the fast timescale *τ*_1_ can be neglected in the second term

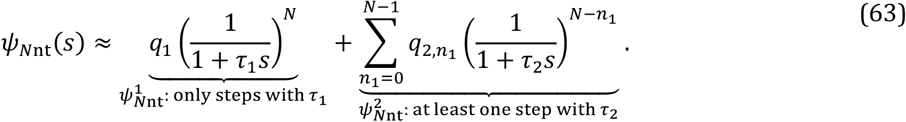

When we take the inverse Laplace transform of this expression, the second term 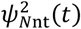 has now been reduced to a sum of single gamma distributions

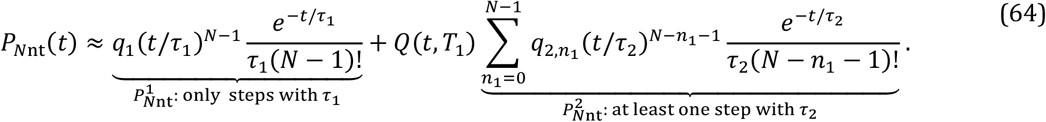

To take into account that the second term is an approximation that only works for long timescales, we added a cut-off factor *Q*(*t, T*_1_) to the term 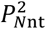 ensuring that the short time behaviour follows that of *N* steps with timescale *τ*_1_ (*T*_1_ = *Nτ*_1_)

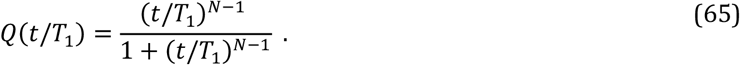

Also, renormalization of the distribution is required, as discussed in the **Main** text.

#### Single exponential approximation for the paths of at least one step with timescale τ_2_

In case a flat shoulder is observed on long timescales in the coarse-grained dwell-time distribution in combination with a peak at short timescales (**Figure 5C**), we can perform a single exponential approximation for the sum of gamma distributions in the second term in **Equation 64** 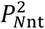. We obtain the overall weight of the shoulder *q*_2_ and average timescale *T*_2_ for all combinations with at least one step with timescale *τ*_2_ using the Laplace transform method in **Equation 52** on the second term in **Equation 60**

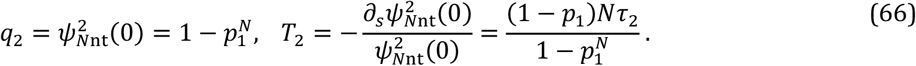

#### Condition on the microscopic rates from the condition on the single step timescales and probabilities

For the model presented in **Figure 5D** we give a condition on the internal rates *k*_f1_ ≫ *k*_f2_ + *k*_I_ + *k*_−I_ which results in the separation of timescales *τ*_1_ ≪ *τ*_2_ and probabilities *p*_1_ ≫ *p*_2_ > 0. In this section, we’ll show how these conditions are derived. First consider that the internal rates are expressed in terms of the single step parameters *τ*_1_, *τ*_2_ and *p*_1_ as in **Equation 34**

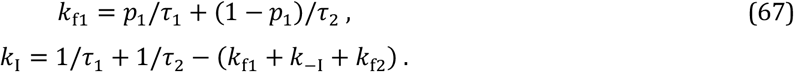

By combining the expressions in **Equation 67**, we obtain *k*_f2_ + *k*_I_ + *k*_−I_ = (1 − *p*_1_)/*τ*_1_ + *p*_1_/*τ*_2_. Because *p*_1_ ≫ *p*_2_ > 0 translates to *p*_1_ ≫ 1/2 and we have *τ*_1_ ≪ *τ*_2_, we conclude that *k*_f1_ ≫ *k*_f2_ + *k*_I_ + *k*_−I_ in this regime.

#### Generalization of the approximate FPT distribution for one narrow peak and multiple flat shoulders

From magnetic tweezers assays on the nucleotide addition cycle of RNA polymerases, we typically obtain coarse-grained dwell-time distributions with a narrow peak at short timescales and more than one flat shoulder on longer timescales (**Figure 1I**) (18, 29, 57). These features can generally be well-fitted with a gamma distribution (**Equation 10**) and exponential distributions with a cut-off at short timescales, respectively

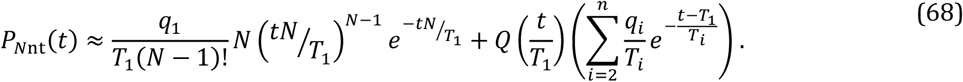

Here we generalized the approximate FPT distribution in **Equation 15** for *n* alternative states with separated timescales *T*_*i*_ and incrementally decreasing weights *q*_*i*_ to satisfy the conditions for flat shoulders (**Main, Figure 5C**).

When these conditions are satisfied, the characteristic timescale *T*_*i*_ and weight *q*_*i*_ of each shoulder can be expressed in terms of the single step parameters *p*_*i*_ and *τ*_*i*_ as

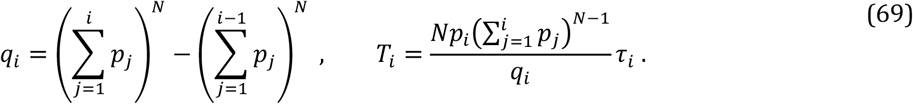

#### Generalization of the FPT distribution for one narrow peak and multiple flat/peaked shoulders

When the coarse-grained dwell-time distribution contains, in addition to a narrow peak at short timescales, multiple well-separated flat or peaked shoulders, each shoulder can in principle be described by a sum of gamma distributions with timescale *τ*_*i*_, such as in **Equation 12** for a single shoulder, provided the corresponding single-step timescales *τ*_*i*_ are sufficiently separated. For example, a distribution consisting of a narrow peak followed by two flat or peaked shoulders at progressively longer timescales can be represented as

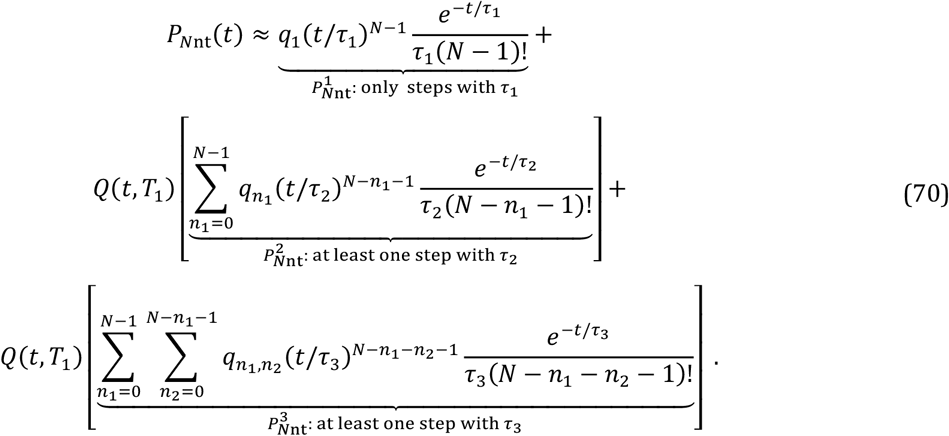

The first term 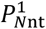 is a gamma distribution with average timescale 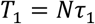 and weight 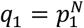, representing the path of *N* steps with timescale *τ*_1_. The second term 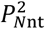 is the sum of gamma distributions with index *n*_1_, average timescale 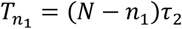, and binomial weight 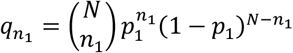, resulting in either a flat or peaked shoulder, which represents the paths with at least one step with timescale *τ*_2_. The third term 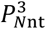 is the sum of gamma distributions with indices *n*_1_ and *n*_2_, average timescale 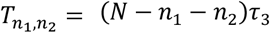 and binomial weight 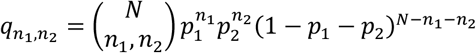, resulting in either a flat or peaked shoulder, which represents the paths with at least one step with timescale *τ*_3_. Both 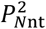 and 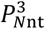 are cut-off at short timescales with *Q*(*t, T*_1_) from **Equation 13** ensuring that the short time behaviour follows that of *N* steps with timescale *τ*_1_. The approximation of the FPT distribution holds for the separation of timescales *τ*_3_ ≫ *τ*_2_ ≫ *τ*_1_.

Also, this distribution needs to be renormalized to the time window of the measurements, as discussed in the **Main** text.

