## Supplementary Information for "Processing, analysing and modelling kinetic data in the era of high-throughput single-molecule biophysics"

**This document includes:**

Figures S1 to S3

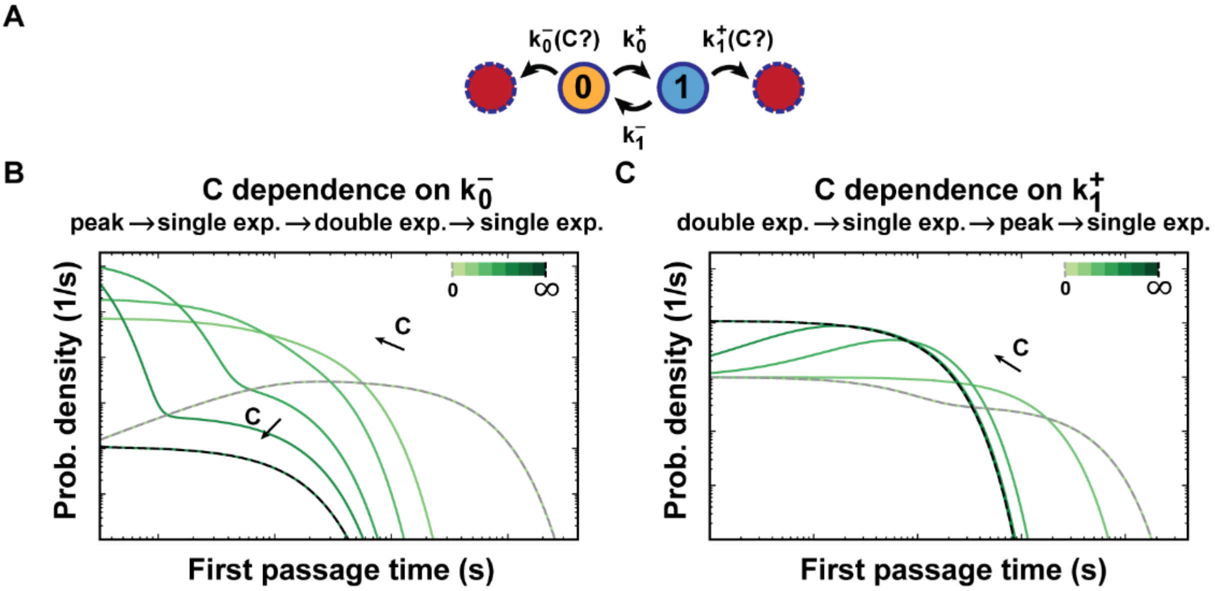


**Figure S1: Dwell-time distributions for two-state systems with a $C$ dependency on an exit rate. (A)** Schematic of two-state systems with a $C$ dependency on one of the exit rates $k_0^-$ or $k_1^+$. **(B)** When $C$ acts on $k_0^-$, the distributions transform from a peak to single to double exponential and back to single exponential with $C$. **(C)** When $C$ acts on $k_1^+$, the distributions transform from double to single exponential to a peak back into a single exponential with $C$.

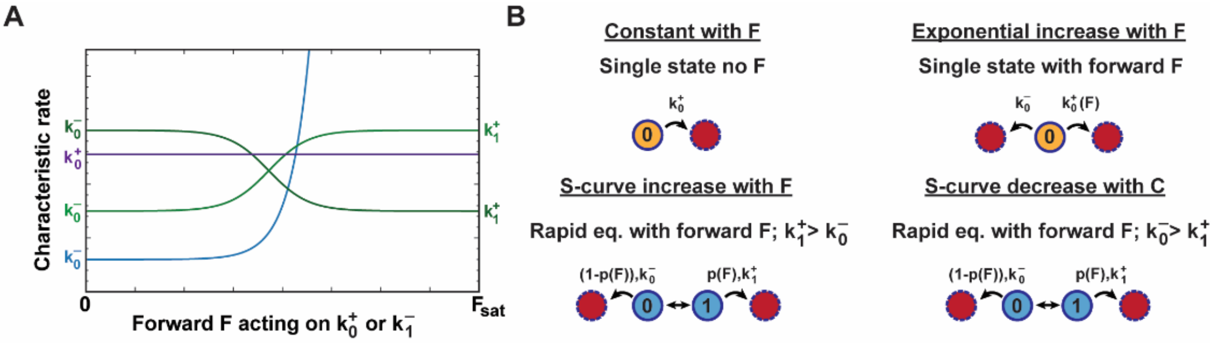


**Figure S2: Minimal cases of single- or two-state systems that show a single shoulder dwell-time distributions with forward force *F* acting on either $k_0^+$ or $k_1^-$. (A)** The typical trends in the characteristic rate with probing force $F$ in the forward (+) direction for single- or two-state systems. The characteristic rate is either constant (purple line), increases exponentially with $F$ (blue line) or shows an S-curve increase or decrease with $F$ (light or dark green line). **(B)** The minimal models in case of single shoulder dwell-time distributions per trend in the characteristic rate with $F$.

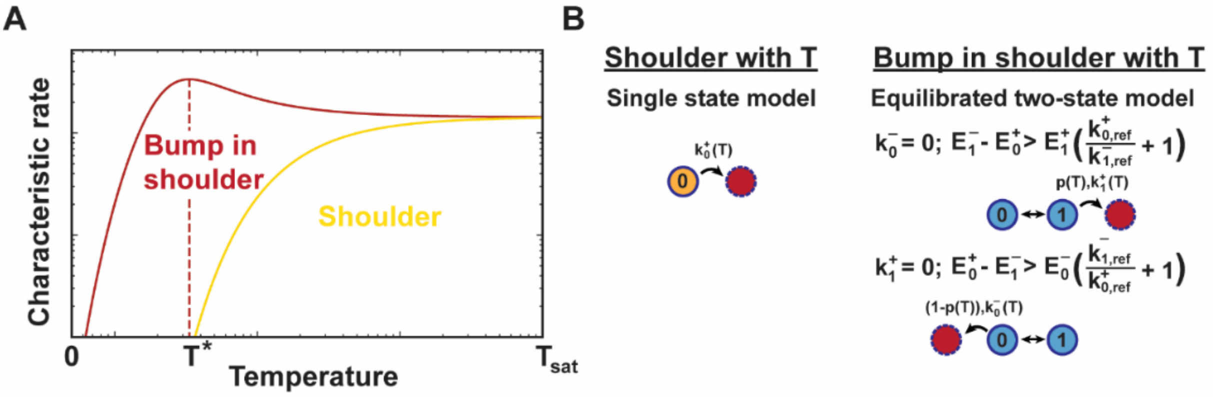


**Figure S3: Distinct trends in the temperature dependency of the characteristic rate for single- and two-state systems with a single exit rate. (A, B)** An intermediate state can be detected from the trend in the characteristic rate with temperature for specific conditions. When the characteristic rate increases monotonically as a "shoulder" with temperature (yellow curve, A) and the minimal model is a single-state system (B). For special conditions on the activation energies of an equilibrated two-state system indicated in panel B, the increase in the characteristic rate at low temperatures changes into an asymptotic decrease for high temperatures, creating a "bump" in the shoulder with the maximum at temperature $T^*$ (red curve, A). For this characteristic trend, the minimal model is an equilibrated two-state system (B).
